# Chemically modified dsRNA induces RNAi effects in insects *in vitro* and *in vivo*: A potential new tool for improving RNA-based plant protection

**DOI:** 10.1101/2022.02.02.478785

**Authors:** John D. Howard, Myriam Beghyn, Nathalie Dewulf, Yves De Vos, Annelies Philips, David Portwood, Peter M. Kilby, Duncan Oliver, Wendy Maddelein, Stephen Brown, Mark J. Dickman

## Abstract

Global agriculture loses over $100 billion of produce annually to crop pests such as insects. Many of these crop pests either have no current means of control or have developed resistance against chemical pesticides. Long dsRNAs are capable of inducing RNA interference (RNAi) in insects and are emerging as novel highly selective alternatives for sustainable insect management strategies. However, there are significant challenges associated with RNAi efficacy in insects. In this study, we synthesised a range of chemically modified long dsRNA in an approach to improve nuclease resistance and RNAi efficacy in insects. The results showed that dsRNA containing phosphorothioate modifications demonstrated increased resistance to southern green stink bug saliva nucleases. Phosphorothioate and 2’-fluoro modified dsRNA also demonstrated increased resistance to degradation by soil nucleases and increased RNAi efficacy in *Drosophila melanogaster* cell cultures. In live insects, chemically modified long dsRNA successfully resulted in mortality in both stink bug and corn rootworm. The results provide further mechanistic insight of RNAi efficacy dependence on modifications in the sense or antisense strand of the dsRNA in insects and demonstrate for the first time that RNAi can successfully be triggered by chemically modified long dsRNA in insect cells or live insects.

## INTRODUCTION

Global agriculture loses over $100 billion of produce annually to crop pests such as insects. Many of these crop pests either have no current means of control or have developed resistance against traditional chemical pesticides. The economic cost of pest damage to agriculture, and particularly the increasing cost due to increasing pesticide resistance is becoming a significant issue (1), with insects consuming 5 to 20% of major grain crops (2). Incidences of pesticide resistance have increased since the 1950s (3), and the spread of the natural range of resistant insects, such as Colorado potato beetle, with climate change threatens to magnify the amount of crop damage done by these insects (4). Just a one degree Celsius rise in temperatures could increase the total losses of rice, corn and wheat alone by 10-25 %, with a two degree Celsius rise resulting in approximately 213 million tons of lost produce (2). In conjunction with an increasing world population, increased loss of produce to pests such as insects also threatens food security, particularly in the developing world.

Another related major driver for the development of new pesticides, is the need for thriving populations of pollinator species such as bees, which are currently being reduced by climate change (5), and Varroa with associated viral infections leading to colony loss (6, 7). There is also a debate about whether the use of existing pesticides results in adverse impacts on populations of beneficial species, for example pollinators such as bees (8).

Beyond the use of traditional small molecule pesticides as a ‘chemical’ method of pest control, an alternative ‘biological’ method of pest control has used *Bacillus thuringiensis* (*Bt*) toxin genes engineered into transgenic crop strains, or *Bt* toxins being applied directly to crops (9). However, many important insect pest species are not susceptible to this method of control, while other previously susceptible species have developed resistance to *Bt* toxins (9). Therefore there is significant demand for the development of new classes of pesticide that can both overcome pesticide resistance of target species by utilising new mechanisms of inducing mortality, and are capable of overcoming issues with lack of target species selectivity, thereby avoiding causing harm to beneficial species.

RNA based biocontrols are emerging as a novel alternative to chemical pesticides for sustainable control of crop pest insects (10–12). RNA based biocontrols, consisting of long dsRNA, are capable of inducing RNA interference (RNAi) in insects, resulting in selective degradation of a target mRNA and therefore reduced levels of its protein product (13). Targeting of the mRNA for a protein essential to the growth and survival of the insect results in mortality of the target insect, therefore, long dsRNA based biocontrols are emerging as novel highly selective insecticides, and have been proposed as a solution to the issues raised above (14).

The first successful studies demonstrating proof-of-principle for this approach in insects took place over ten years ago (10, 11), and in the intervening years research has been undertaken demonstrating the possibility of using this approach for a wide range of targets and insect species (15–20). In many species, triggering RNAi has been demonstrated to be highly effective, inducing poor health and mortality of the target insects fast enough to significantly protect crop plants (17).

However, there are differences in RNAi efficacy between different insect orders and species due to variation in factors such as insect nuclease potency and upregulation (21), physiological pH (22), and dsRNA uptake and subsequent intracellular transport (22). Long dsRNA based insecticides can also be degraded by nucleases, either in the environment (e.g. in soil) (23), or in the bodily fluids of the target insect (24). Some insect orders such as Lepidoptera demonstrate greater nuclease degradation of dsRNA *in vivo* than others (21). Degradation of dsRNA by nucleases in the environment and within the insect’s body before it can induce RNAi is a major barrier to successful triggering of RNAi by ingestion of dsRNA in insects (24, 25). Resistance of dsRNA to degrading nucleases, and successful processing of dsRNA by the insect Dicer-2 nuclease – a key component of the insect RNAi pathway (26) – are therefore key factors that also affect the efficacy of dsRNA based biocontrols.

The application of RNA based products for insect management strategies, typically requires long dsRNA substrates of at least 50 bp, with only dsRNAs of over 100 or 200 bp being effective in some insect species (17, 27, 28). In contrast, therapeutic short interfering RNAs (siRNAs) and DNA antisense oligonucleotides are between 8 and 50 bp in length (29) and require a range of chemical modifications and appropriate formulations to ensure efficacy in whole organism mammalian systems (29, 30). The chemical modifications prevent their degradation in the bloodstream by extracellular nucleases (31), as well as improving delivery and transport of the siRNA (30).

Research into the use of RNAi as a therapeutic method has seen a large number of different types of RNA chemical modifications investigated, as reviewed in Shen and Corey (30). Modifications to the bases themselves have seen some investigation (32), though most attention has focused on modifications to the ribose-phosphate backbone. Phosphorothioate modifications have been the most commonly utilised, as have modifications to the 2’ position of the ribose sugar ring including 2’-F and 2’-O-Me, and locked and unlocked nucleic acid (LNA and UNA) modifications have also been used (33–37).

Several of the chemical modifications examined here are known to increase resistance of siRNAs (38, 39), antisense ss-siRNAs (39), antisense oligonucleotides (40), and chimeric oligonucleotides (41) to nuclease degradation in mammalian or other systems. Therefore, it was proposed that including these chemical modifications in long dsRNA could provide increased resistance to insect nucleases compared to unmodified dsRNA. This protection could potentially improve RNAi efficacy compared to unmodified dsRNA, thus reducing the dose of dsRNA required to achieve high mortality of a target pest insect on a crop.

In this study, we have optimised the synthesis and purification of long dsRNA containing a range of different chemical modifications including phosphorothioate (PS), 2’-fluoro (2’F), and 5-hydroxymethyl (HMr) modifications (see Table 1). The effects of chemical modifications on the nuclease stability of long dsRNA were studied *in vitro* using southern green stink bug (*Nezara viridula*) (SGSB) saliva, Colorado potato beetle (*Leptinotarsa decemlineata*) (CPB) gut secretions, and agricultural soil as examples of sources of nucleases likely to contribute to degradation of insecticidal dsRNA. The ability of model RNase III/Dicer family enzymes to successfully cleave long chemically modified dsRNA into endoribonucleace-prepared siRNAs (esiRNAs) *in vitro* was also investigated. Finally, the RNAi efficacy of long chemically modified dsRNA was examined both *in vitro* in *Drosophila* cell culture using a dual luciferase reporter assay, and *in vivo* in SGSB nymphs and western corn rootworm (*Diabrotica virgifera virgifera*) (WCR) larvae using survival studies.

**Table 1.** Composition and nomenclature of chemically modified dsRNA synthesised and used in this study.

| **dsRNA** | ***In vitro* transcription** | **Resulting dsRNA** | **Pooled** | **Name** |
| --- | --- | --- | --- | --- |
| Phosphorothioate dsRNA | Replacement of NTPs with α-thiophosphate ATP or CTP or GTP or UTP | Phosphorothioate (A)<br>Phosphorothioate (C)<br>Phosphorothioate (G)<br>Phosphorothioate (U) | All 4 | 1PS |
| Phosphorothioate dsRNA | Replacement of NTPs with combinations of α-thiophosphates ATP/CTP/GTP/UTP | Phosphorothioate (AC)<br>Phosphorothioate (AG)<br>Phosphorothioate (AU)<br>Phosphorothioate (CG)<br>Phosphorothioate (CU)<br>Phosphorothioate (GU) | All 6 | 2PS |
| 2'-Fluoro dsRNA | Replacement of CTP or UTP with 2'-Fluoro CTP or UTP | 2'-Fluoro (C)<br>2'-Fluoro (U) | All 2 | 1 2'F |
| 2'-Fluoro dsRNA | Replacement of CTP/UTP with 2'-Fluoro CTP and UTP | 2'Fluoro (CU) | Single | 2 2'F |
| Hydroxymethyl dsRNA | Replacement of CTP or UTP with 5-Hydroxymethyl CTP or UTP | Hydroxymethyl (C)<br>Hydroxymethyl (U) | All 2 | 1HMr |
| Hydroxymethyl dsRNA | Replacement of CTP/UTP with 5-Hydroxymethyl CTP and UTP | Hydroxymethyl (CU) | Single | 2HMr |

The results showed for the first time that long dsRNA containing phosphorothioate modifications demonstrated increased resistance to stink bug saliva nucleases. In addition, both phosphorothioate and 2’-fluoro modified dsRNAs demonstrated increased resistance to soil nuclease degradation, and increased RNAi efficacy in *Drosophila* cell cultures. Furthermore, the effects of the chemical modifications of long dsRNA on RNAi efficacy were also studied in live insects in both SGSB using injection assays, and in western corn rootworm using feeding assays. The results demonstrate that the chemically modified long dsRNA resulted in successful RNAi in live insects as measured by insect mortality. To our knowledge this is the first time that RNAi has successfully been triggered by chemically modified long dsRNA in insect cells or live insects. Differences in RNAi efficacy *in vivo* were also observed depending on whether modifications were in the antisense strand (i.e. the intended guide strand), the sense strand (i.e. the intended passenger strand), or both strands.

These results provide further mechanistic insight into the effects of chemical modifications of dsRNA used in plant protection. It is anticipated that these results will provide important information for developing new alternative dsRNA based plant protection products with improved nuclease resistance and RNAi efficacy.

## RESULTS

### Synthesis of chemically modified dsRNA by *in vitro* transcription (IVT) and analysis using gel electrophoresis and ion pair reverse phase (IP RP) HPLC

A range of unmodified and chemically modified dsRNA were synthesised from DNA templates by *in vitro* transcription (IVT). Chemically modified RNA was synthesised by substituting canonical natural nucleotide triphosphates (NTPs) for chemically modified NTP analogues in the IVT reaction, generating RNA with all of one or two of the four canonical nucleotides replaced by chemically modified analogues in either the sense, antisense, or both strands (see Figure 1A for a description of the nomenclature used). T7 RNA polymerase was used in IVT reactions to synthesise unmodified, phosphorothioate modified, and 5-hydroxymethyl modified RNA. T7R&DNA polymerase was used in IVT reactions to synthesise 2’-fluoro modified RNA. Structures of the modified nucleotides incorporated are shown in Figure 1B.

**Figure 1.**
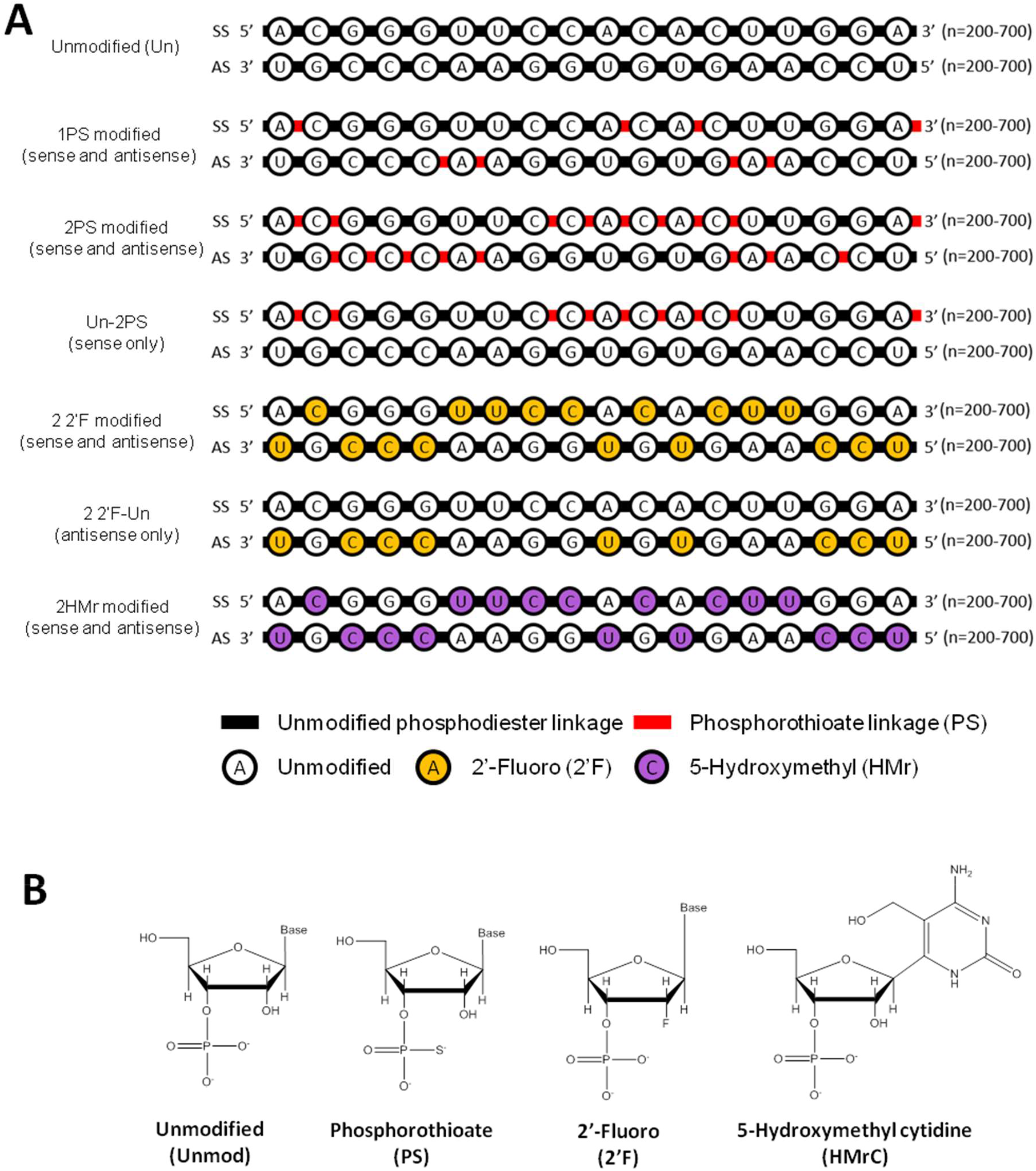
Nomenclature for chemically modified dsRNA and structure of chemically modified nucleotides. (**A**) Illustration of nomenclature for phosphorothioate (PS), 2’-fluoro (2’F) and 5-hydroxymethyl (HMr) modified dsRNA (typically 200-700 bp). dsRNA with phosphorothioate linkages (PS), were synthesised in IVT reactions with one canonical NTP replaced by an α-thio NTP analogue and are referred to as 1PS. dsRNAs synthesised in IVT reactions with two NTPs replaced by their corresponding α-thio NTP analogues are referred to as 2PS. dsRNAs with modifications in one strand are referred to as “Un-2PS”, “2PS-Un” with the format “antisense strand-sense strand”, or “guide strand-passenger strand” in terms of functional RISCs (discounting non-targeting RISCs where the sense strand is loaded as the guide strand). Nomenclature for dsRNAs with 2’F and HMr modifications is the same as for PS, as per the examples shown, and these dsRNAs were also synthesised in IVT reactions, with one or more canonical NTP replaced by a 2’-fluoro NTP or 5-hydroxymethyl NTP analogue. (**B**) Structures of the modified nucleotide analogues.

Purified dsRNA was analysed by gel electrophoresis and ion pair reverse phase HPLC to validate the synthesis of the chemically modified dsRNA and confirm the purity of the RNA sample (Supplementary Figures S1&2). Quantification of the dsRNA was performed using UV spectrophotometry and confirmed using the HPLC peak areas to ensure accurate quantification of the dsRNA prior to downstream applications. For use in functional assays, dsRNA containing one or more modified nucleotide analogues were in many cases mixed into pooled samples, for example – 2’F C dsRNA and 2’F U dsRNA were combined into a mixed pool of 1 2’F dsRNA. These pooled samples were as described in Table 1.

### *In vitro* investigation of the effect of dsRNA chemical modifications on dsRNA degradation by nucleases and UV exposure

dsRNA biocontrols applied in the field face many sources of degradation prior to uptake by the cells of the target insect. dsRNA applied to leaf surfaces may be degraded by UV exposure, whereas dsRNA applied to soil, to target soil-dwelling insect crop pests may be degraded by the activity of nucleases present in the soil. Once ingested by the target insect, dsRNA has to pass through bodily fluids such as saliva, gut secretions, and hemolymph before reaching the RNAi machinery inside individual insect cells, and all of these fluids contain nucleases which may degrade the dsRNA en route, with the nuclease activity of these bodily fluids varying greatly between different insect species.

In order to investigate the effects of chemical modifications on the resistance of dsRNA to degradation by insect and environmental nucleases, and other environmental factors, unmodified and chemically modified dsRNA were exposed to a variety of different model nucleases/extracts and the degradation of unmodified and chemically modified dsRNA was compared. The model nucleases and environmental factors selected for testing were as follows. SGSB saliva was selected as it contains high nuclease activity, which may degrade RNA and therefore negatively affect the use of RNA based biocontrols (42). In addition, CPB gut secretions, which contain both insect and plant nucleases were selected. Furthermore, CPB is a highly relevant pest species resistant to many small molecule pesticides but susceptible to RNA based biocontrols (17). Agricultural soil effluent was also used as nuclease degradation of dsRNA in soil is a possible barrier to successful insecticidal RNAi (23) especially for soil dwelling crop pests such as corn rootworm. Finally, the effect of UV radiation exposure to dsRNA was studied, as UV radiation has previously been shown to degrade dsRNA (17). In each assay, differences in degradation between unmodified and chemically modified dsRNA were determined using gel electrophoresis. The dsRNA band intensities for each replicate of each time point and/or dilution were normalised against the corresponding band intensity of the zero time point or water only control to calculate the relative dsRNA stability index.

In order to determine potential differences in nuclease resistance between unmodified and chemically modified dsRNA, a number of chemically modified dsRNAs were incubated with a range of dilutions of SGSB saliva-containing Sf9 media, and samples removed for analysis by gel electrophoresis at a number of time points (see Figure 2A). The band intensities for each replicate of each time point and media dilution were normalised against the corresponding band intensity of the water only control band, to calculate the “relative dsRNA stability” (see Figure 2B and Supplementary Figure S3). The results demonstrate that phosphorothioate dsRNA was more resistant to stink bug saliva nuclease degradation than unmodified dsRNA. In contrast, no increase in stability of the 2’F or 5-hydroxymethyl modified dsRNA was observed. The differences are most pronounced at the 1/27 and 1/81 dilutions of saliva-containing media and examples of these are highlighted in red boxes in Figure 2A. dsRNA with PS or 2’F modifications in only one strand demonstrated intermediate stability between that of unmodified dsRNA and dsRNA with the same chemical modification present in both strands (see Supplementary Figure S3).

**Figure 2.**
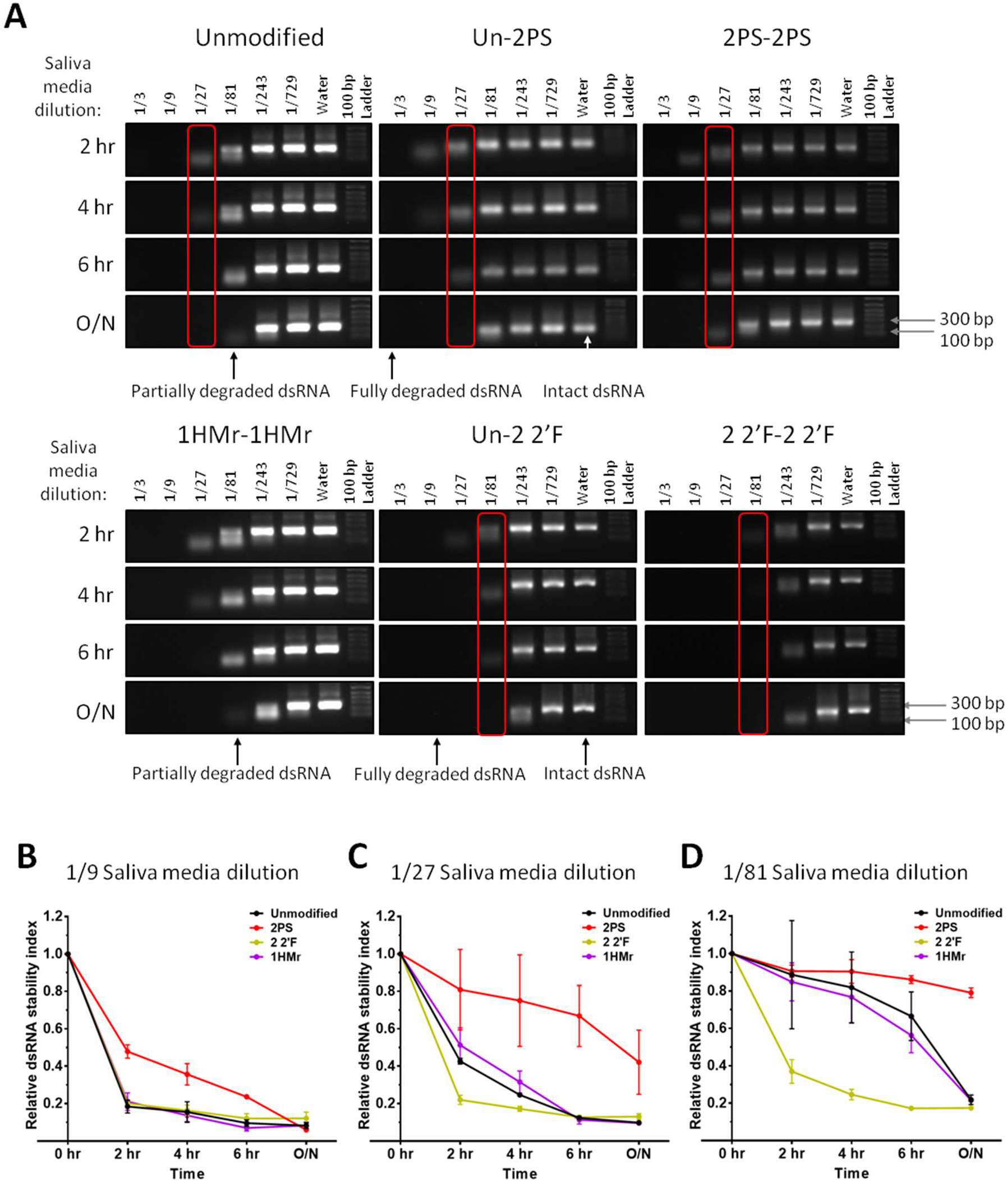
Nuclease stability assays to determine the resistance of chemically modified dsRNA to degradation by stink bug saliva nucleases. (**A**) Gel electrophoretograms of unmodified, Un-2PS, 2PS-2PS, Un-2 2’F, 2 2’F-2 2’F, and 1HMr-1HMr Target B dsRNA incubated at room temperature with Sf9 cell culture medium containing SGSB saliva. Samples were incubated with saliva contaminated medium at a range of dilutions in water and reactions stopped at either 2 hrs, 4 hrs, 6 hrs or after incubation overnight (O/N) by addition of formamide loading dye and freezing at −20 °C. (**B-C**) Time course graphs showing quantification of gel band intensity from (**A**) with the average and SD of *n* = 2 replicate sets of incubations and gels plotted for a subset of key saliva medium dilutions, 1/9 dilution (**B**), 1/27 dilution (**C**), and 1/81 dilution (**D**).

Chemically modified dsRNA was also incubated in a range of dilutions of CPB gut secretions prior to analysis by gel electrophoresis and the band intensities used to determine the relative dsRNA stability (Supplementary Figure S4 A-C). The results show that in contrast to the stink bug saliva nuclease assay, there was no difference in stability of the dsRNA to degradation by CPB gut nucleases at any concentration, between unmodified, phosphorothioate modified, and 2’-F modified dsRNA. Unmodified and phosphorothioate dsRNA incubated with a 1/1,000 dilution of CPB gut secretions for a time course assay also demonstrated no difference in stability at any time point (Supplementary Figure S4 D&E). Further analysis was performed by incubating chemically modified dsRNA with an aqueous extract from agricultural soil prior to analysis by gel electrophoresis (see Figure 3A) and the band intensities used to determine the relative dsRNA stability (see Figure 3B). The results demonstrate that both PS and 2’F modified dsRNA are more resistant to degradation by soil nucleases compared to unmodified dsRNA.

**Figure 3.**
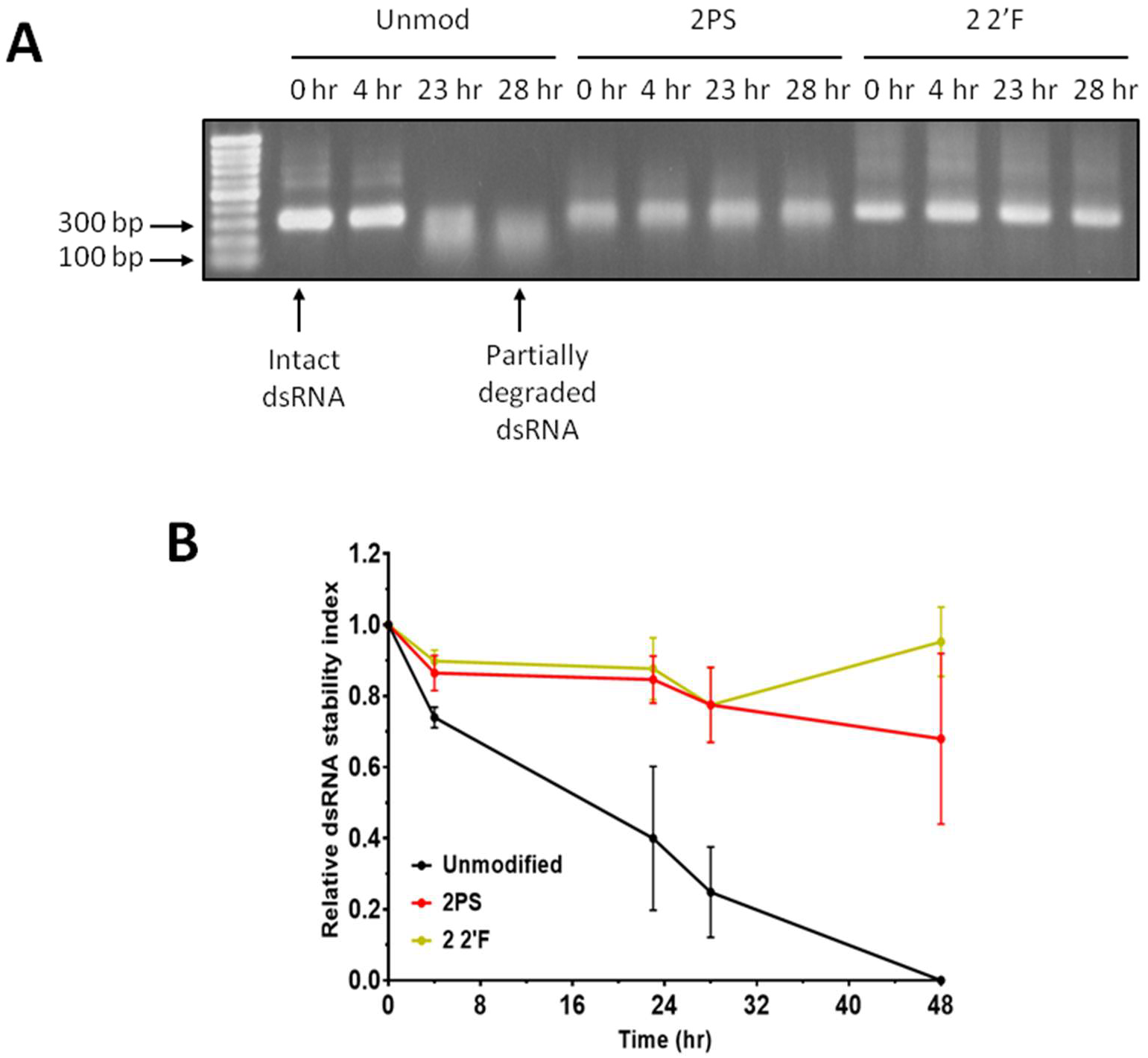
Nuclease stability assays to determine the resistance of chemically modified dsRNA to degradation by agricultural soil nucleases. (**A**) Gel electrophoretograms of unmodified, 2PS-2PS, and 2 2’F-2 2’F Target B dsRNA incubated at 37 °C with water containing soil nucleases. Reactions stopped at a range of time points by addition of formamide loading dye and freezing at −20 °C. (**B**) Time course graph showing quantification of gel band intensity from (**A**) and further replicates with the average and SD plotted. 48 hrs: *n* = 2, Other time points: *n* = 3.

Degradation of dsRNA by UV radiation was also assessed in a similar manner to (17). Samples were taken at a number of time points, and analysed by gel electrophoresis and the band intensities used to determine the relative dsRNA stability (Supplementary Figure S4F&G). The results demonstrate that there was no difference in resistance of the dsRNA to UV degradation between unmodified and phosphorothioate modified dsRNA. The results for unmodified dsRNA are consistent with previous studies that demonstrated significant dsRNA degradation was observed after 2 hrs UV exposure (17). The results demonstrate clear differences in stability between unmodified and chemically modified dsRNAs for both the saliva and soil nuclease assays. In contrast, the rate of dsRNA degradation by CPB gut secretions or UV radiation was equal for unmodified or chemically modified dsRNA. These results highlight that different dsRNA chemical modifications potentially need to be utilised dependent upon the major source of dsRNA degradation in a particular application environment.

### Processing of chemically modified dsRNA substrates *in vitro* by Dicer/RNase III enzymes

For insecticidal dsRNA to be functional in insect cells, they must be capable of being processed into functional siRNAs by the Dicer-2 enzyme (43). This processing may be affected by the presence of chemical modifications in the dsRNA, either at the binding step or cleavage step. In order to examine the ability of Dicer enzymes to process chemically modified long dsRNA into endoribonuclease-prepared siRNAs (esiRNAs), two different types of Dicer/RNase III family enzyme were used: bacterial RNase III, and *Giardia intestinalis* Dicer. These two enzymes bind and process dsRNA in distinct ways and contain fewer functional domains than insect or mammalian Dicer enzymes (see Supplementary Figure S5). By studying the activity of bacterial RNase III and *Giardia intestinalis* Dicer on chemically modified dsRNA *in vitro*, it was proposed that this would provide further insight into the potential effect of such chemical modifications on insect Dicer-2 dsRNA processing *in vivo*.

*In vitro* assays were performed using unmodified, phosphorothioate, 2’-fluoro and 5-hydroxymethyl dsRNA to determine the efficacy of processing of the chemically modified dsRNA to esiRNAs by both enzymes. DsRNA was incubated with RNase III or *Giardia* Dicer, followed by analysis by gel electrophoresis (see Figure 4). The results show that bacterial RNase III successfully cleaved all the unmodified and chemically modified dsRNA substrates to esiRNAs (Figure 4 A&B). *Giardia* Dicer successfully generated esiRNAs from unmodified dsRNA, however failed to fully process 1HMr, 2PS and 2 2’F dsRNA substrates into esiRNA products (Figure 4C&D). Further analysis of phosphorothioate dsRNA demonstrated the number of phosphorothioate linkages in the dsRNA correlated with increased resistance towards *Giardia* Dicer processing the dsRNA substrate to esiRNAs (see Figure 4C). dsRNA with 2’-fluoro modifications in only one strand also demonstrated greater suitability as a substrate for *Giardia* Dicer than dsRNA with 2’-fluoro modifications in both strands (Figure 4D). However, dsRNA with 2’-fluoro modifications in both strands was still partly processed to generate esIRNAs. While 2 2’F dsRNA was not completely processed by *Giardia* Dicer *in vitro* under the conditions used, there is evidence from previous studies that indicate that 1 2’F dsRNA with a lower number of 2’-fluoro modifications can be effectively processed by *Giardia* Dicer (44).

**Figure 4.**
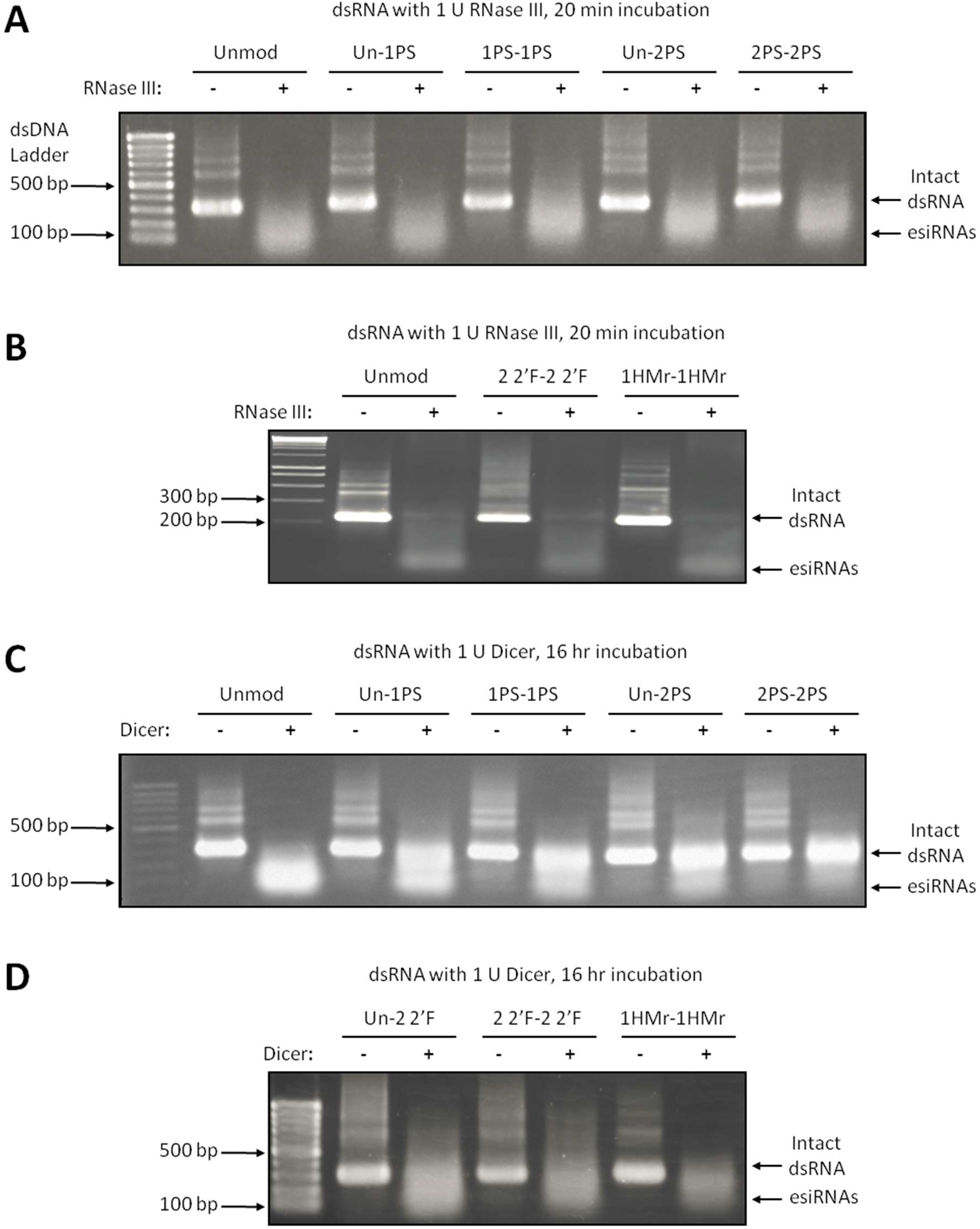
Processing of unmodified and chemically modified dsRNA to esiRNAs by Dicer/RNase III family enzymes (*E. coli* RNase III and *Giardia intestinalis* PowerCut model Dicer). (**A, B**) Processing of unmodified and chemically modified dsRNA to esiRNAs by bacterial RNase III. (**A**) Gel electrophoretogram of unmodified, Un-1PS, 1PS, Un-2PS, and 2PS Target B dsRNA incubated with 1 U of RNase III for 20 mins. (**B**) Gel electrophoretogram of unmodified, 2 2’F and 1HMr Target B dsRNA incubated with 1 U of RNase III for 20 mins. (**C, D**) Processing of unmodified and chemically modified dsRNA to esiRNAs by *Giardia* ‘PowerCut’ Dicer. (**C**) Gel electrophoretogram of unmodified, Un-1PS, 1PS, Un-2PS, and 2PS Target B dsRNA incubated with 1 U *Giardia* Dicer for 16 hr. (**D**) Gel electrophoretogram of Un-2 2’F, 2 2’F, and 1HMr Target B dsRNA incubated with 1 U *Giardia* Dicer for 16 hr. All incubations were performed at 37 °C; 1 μg of dsRNA was incubated with 1 U RNase III or PowerCut Dicer; reactions were quenched by addition of EDTA. In each gel a dsDNA ladder, intact dsRNA and processed esiRNAs are highlighted.

### Chemically modified dsRNA induces RNAi *in vitro* in an insect cell line (*Drosophila* Kc167)

Our first aim was to establish whether chemically modified dsRNA could induce RNAi in insect cells, and to study the effects of dsRNA chemical modifications on RNAi efficacy. *Drosophila melanogaster* Kc167 cells were selected as the model system in which to assess these factors. Insects that exhibit RNAi from the ingestion of dsRNA, uptake the dsRNA from their gut lumen by receptor-mediated endocytosis (27, 45, 46). *Drosophila* cells in culture similarly uptake naked dsRNA via scavenger receptors such as Eater, which bind the dsRNA and initiate its uptake by receptor mediated endocytosis (27, 47). As no transfection reagent is used for delivery of the dsRNA into the cells, the ability of a transfection reagent to bind and facilitate the uptake of modified dsRNA is not a factor in the efficacy of the dsRNA. Therefore, this represents a realistic model of the uptake of, and RNAi-induced mRNA degradation by, dsRNA in the cells of live insects. In order to ascertain changes in RNAi efficacy due to the presence of chemical modifications, the ability of the dsRNA to trigger degradation of their target mRNA was quantified by a dual luciferase assay reporter system.

Quantitative analysis of RNAi-induced degradation was carried out with luciferase assays in 96 well plates containing Kc167 cells transfected with luciferase reporter system plasmids. A wide range of dsRNA concentrations (1,000 ng to 0.01 ng per well) were used in order to generate dose curves for unmodified, 1PS, 2PS, 1 2’F, 2 2’F, 1HMr, and 2HMr FLuc dsRNA. Unmodified F59C6.5 non-targeting dsRNA at 1000 ng per well was used as a control, along with controls with no dsRNA. Cells transfected with the luciferase reporter plasmids were incubated with dsRNA for four days, prior to lysis, addition of the luciferase assay reagents, and luciferase luminescence quantification using a plate reader. FL/RL values for wells containing dsRNA were normalised to the mean FL/RL value of wells containing no dsRNA.

Analysis of control luciferase luminescence (RL) values normalised to the mean RL value of wells containing no dsRNA, demonstrated all normalise RL values for all concentrations of all unmodified and chemically modified dsRNA were close to 1 (Supplementary Figure S6) and therefore none of the chemically modified dsRNA were cytotoxic. The three dose curves of normalised FL/RL values for unmodified FLuc dsRNA from the three assays assessing PS, 2’-F and HMr FLuc dsRNA were compared, and demonstrated that duplicate results generated at the same time for the same dsRNA were highly reproducible (Supplementary Figure S7). Together these results give confidence that differences in normalised FL/RL values between unmodified FLuc dsRNA and chemically modified FLuc dsRNA, are due to differences in RNAi efficacy between chemically modified and unmodified dsRNA, and not a result of cytotoxic effects or assay variation.

The results of the luciferase assays comparing RNAi efficacy of unmodified and chemically modified FLuc dsRNA are shown in Figure 5 (complete replicate data Supplementary Figure S8). The results of the luciferase assay comparing RNAi efficacy of unmodified and phosphorothioate (PS) FLuc dsRNA, demonstrate that at the highest concentration of 1 μg of dsRNA per well unmodified, 1PS, and 2PS FLuc dsRNAs were equally effective at inducing firefly luciferase reductiion compared to non-targeting F59C6.5 control dsRNA (Figure 5B). The results of the luciferase assay comparing RNAi efficacy of unmodified and phosphorothioate (PS) FLuc dsRNA, demonstrate that at 10 ng (Figure 5A) 2PS dsRNA was significantly more efficacious than unmodified dsRNA. The full dose curves from the assay (Figure 5C) were used to determine IC50 values which show that the 1PS dsRNA (IC50: 7.6 ng) and 2PS dsRNA (IC50: 2.8 ng) were lower than unmodified dsRNA (IC50: 20.6 ng) consistent with the previous observations.

**Figure 5.**
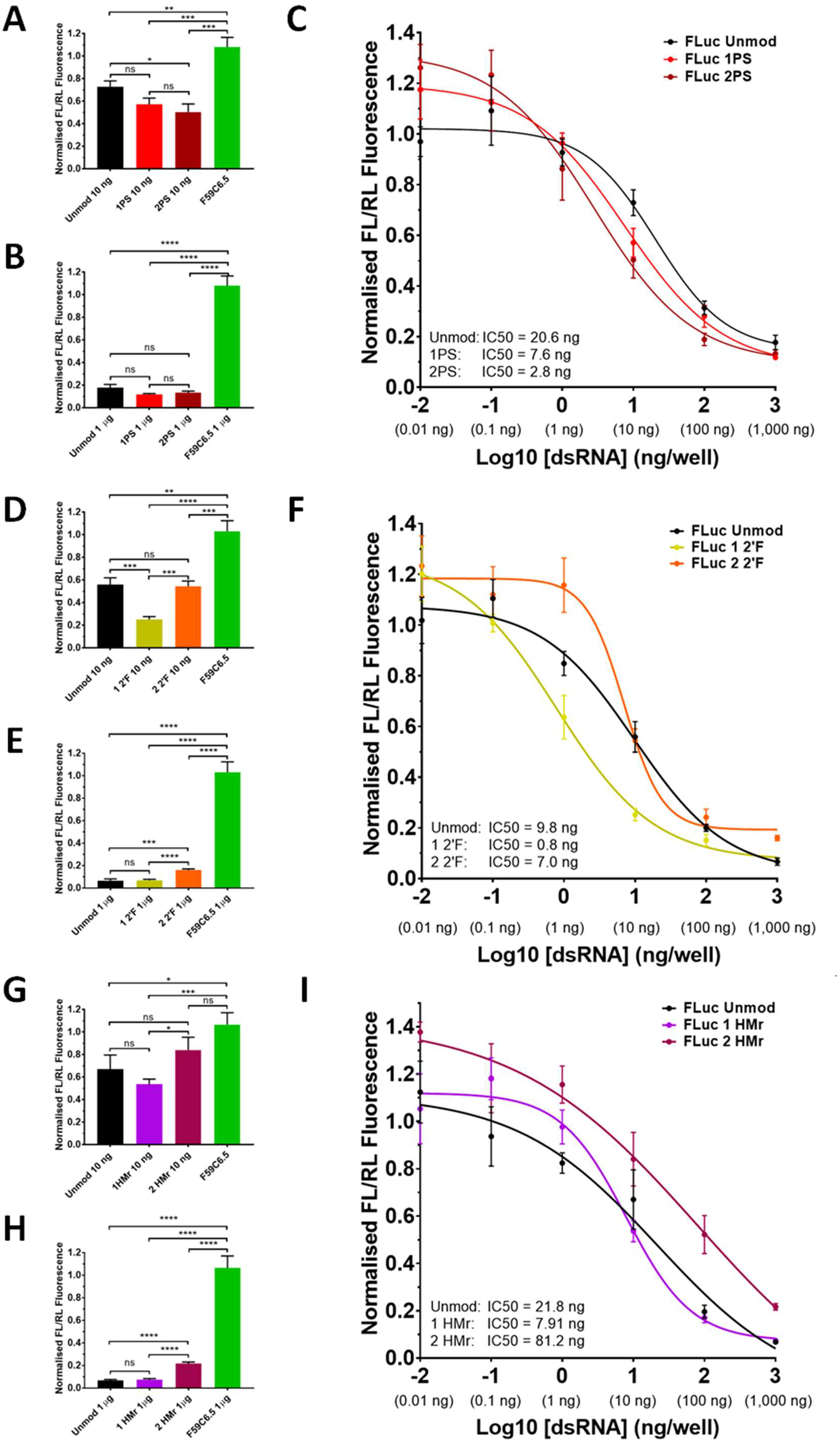
*In vitro* analysis of the effects of dsRNA chemical modifications on RNAi in insect cells across a range of dsRNA concentrations. Results showing quantification of RNAi effects on a firefly luciferase reporter in *Drosophila* Kc167 cell cultures, quantified using a dual luciferase assay reporter system. RNAi effect on a firefly (*Photinus pyralis*) luciferase reporter by FLuc dsRNA, is presented as ratios (FL/RL) of firefly luciferase luminescence intensity (FL) to control sea pansy (*Renilla reniformis*) luciferase luminescence intensity (RL), normalised against FL/RL values for control conditions with no dsRNA. (**A-C**) Results for phosphorothioate (PS) and unmodified (Unmod) FLuc dsRNA. (**D-F**) Results for 2’-fluoro (2’F) and unmodified (Unmod) FLuc dsRNA. (**G-H**) Results for 5-hydroxymethyl (HMr) and unmodified (Unmod) FLuc dsRNA. (**A, D, G**) Bar graphs of RNAi knockdown by 10 ng of Unmod, 1 modified and 2 modified FLuc dsRNA per well against F59C6.5 control dsRNA. (**B, E, H**) Bar graphs of RNAi effect by 1 μg of Unmod, 1 modified and 2 modified FLuc dsRNA per well against F59C6.5 control dsRNA. (**C, F, I**) Dose curves of normalised FL/RL values plotted against log of dsRNA dose per well in ng. EC50s are concentrations at which 50% reductionoccurs. Coloured stars denote statistical significance of the identically coloured data point compared to unmodified dsRNA. Curves generated by non-linear regression analysis using a dose-response inhibition variable slope model in Graphpad prism software. In all graphs, mean and SEM are plotted of *n* = 6 technical replicates. Stars denote significance as calculated by unpaired T-tests. ns = P > 0.05, * = P ≤ 0.05, ** = P ≤ 0.01, *** = P ≤ 0.001, **** = P ≤ 0.0001.

The results of the luciferase assay comparing RNAi efficacy of unmodified and 2’-fluoro (2’F) FLuc dsRNA are shown in Figure 5 D-F. The results demonstrate that at the highest concentration of 1 μg of dsRNA per well unmodified and 1 2’F FLuc dsRNAs were equally effective at inducing firefly luciferase reduction(90-95%) compared to non-targeting F59C6.5 control dsRNA. However, 2 2’F FLuc dsRNA demonstrated reduced RNAi efficacy at this concentration with an RNAi-induced reduction efficiency of approximately 85% (Figure 5D). At 10 ng of dsRNA, 1 2’F dsRNA was significantly more efficacious than unmodified dsRNA, whereas the efficacy of 2 2’F dsRNA at this concentration was the same as unmodified dsRNA (Figure 5C). The full dose curves from the assay (Figure 5E) were used to determine IC50 values and show that 1 2’F dsRNA demonstrates an improvement in RNAi efficacy with an IC50 (0.8 ng) compared to unmodified dsRNA IC50 (9.8 ng). 2 2’F dsRNA demonstrated a similar IC50 (7.0 ng) to unmodified dsRNA, consistent with previous analysis.

The results of the luciferase assay comparing RNAi efficacy of unmodified and 5-hydroxymethyl (HMr) FLuc dsRNA are shown in Figure 5G-I. The results show that at the highest concentration of 1 μg of dsRNA per well unmodified and 1HMr FLuc dsRNAs were equally effective at inducing firefly luciferase reduction(90-95%) compared to non-targeting F59C6.5 control dsRNA. However, 2HMr FLuc dsRNA demonstrated reduced RNAi efficacy at this concentration with an RNAi-induced reduction of approximately 80% (Figure 5H), significantly less than for the same concentration of unmodified dsRNA. At 10 ng of dsRNA, the efficacies of 1HMr and 2HMr dsRNA were not statistically significant from that of unmodified dsRNA (Figure 5G). The full dose curves from the assay (Figure 5I) were used to determine IC50 values and show that the lowest IC50 value was obtained for 1HMr dsRNA (7.9 ng) compared to unmodified dsRNA (21.8 ng). In contrast, the IC50 value of 2HMr dsRNA was 81.2 ng indicating a reduction in RNAi efficacy consistent with previous observations.

### Chemically modified dsRNA can induce RNAi resulting in mortality, *in vivo* in live stink bug (*Nezara viridula*) when delivered by microinjection

For the successful application of chemically modified dsRNA insect control, they must successfully induce RNAi of a target mRNA in the target insect, and RNAi induced degradation of the target mRNA must result in the mortality of the insect. Conversely, the dsRNA must not induce mortality through toxicity of the chemical modifications, in order to preserve the high selectivity of the method. In addition, the chemically modified dsRNA must be successfully taken up when delivered orally to insects through a mechanism which allows it to induce RNAi once inside individual cells. A chemically modified dsRNA should fulfil these criteria in order to be an effective RNA based insect control.

In order to determine whether chemically modified dsRNA could induce RNAi in insects when directly delivered, chemically modified dsRNA was injected into N2 stage SGSB nymphs, and their survival monitored over the course of a number of days. Stink bugs were chosen as a model organism as they are a highly relevant potential target crop pest insect for a dsRNA insecticide product, and their size is convenient for both ease of injection and the dose of dsRNA required. An initial assay in which a large amount of dsRNA was delivered to the insects, demonstrated that unmodified dsRNA and dsRNA containing phosphorothioate (2PS) or 2’-fluoro (2 2’F) modifications in either the sense or antisense strand could induce high levels of mortality through RNAi of a critical target mRNA compared to control injections of water or a non-targeting GFP dsRNA (Figure 6A). Further analysis was performed, in which precise and controlled doses of dsRNA were delivered into N2 stink bugs confirmed that both unmodified and phosphorothioate (1PS in both strands) dsRNA could induce high levels of mortality, and demonstrated that non-targeting control dsRNA containing phosphorothioate modifications (2PS in both strands) did not induce mortality through toxic effects (Figure 6B).

**Figure 6.**
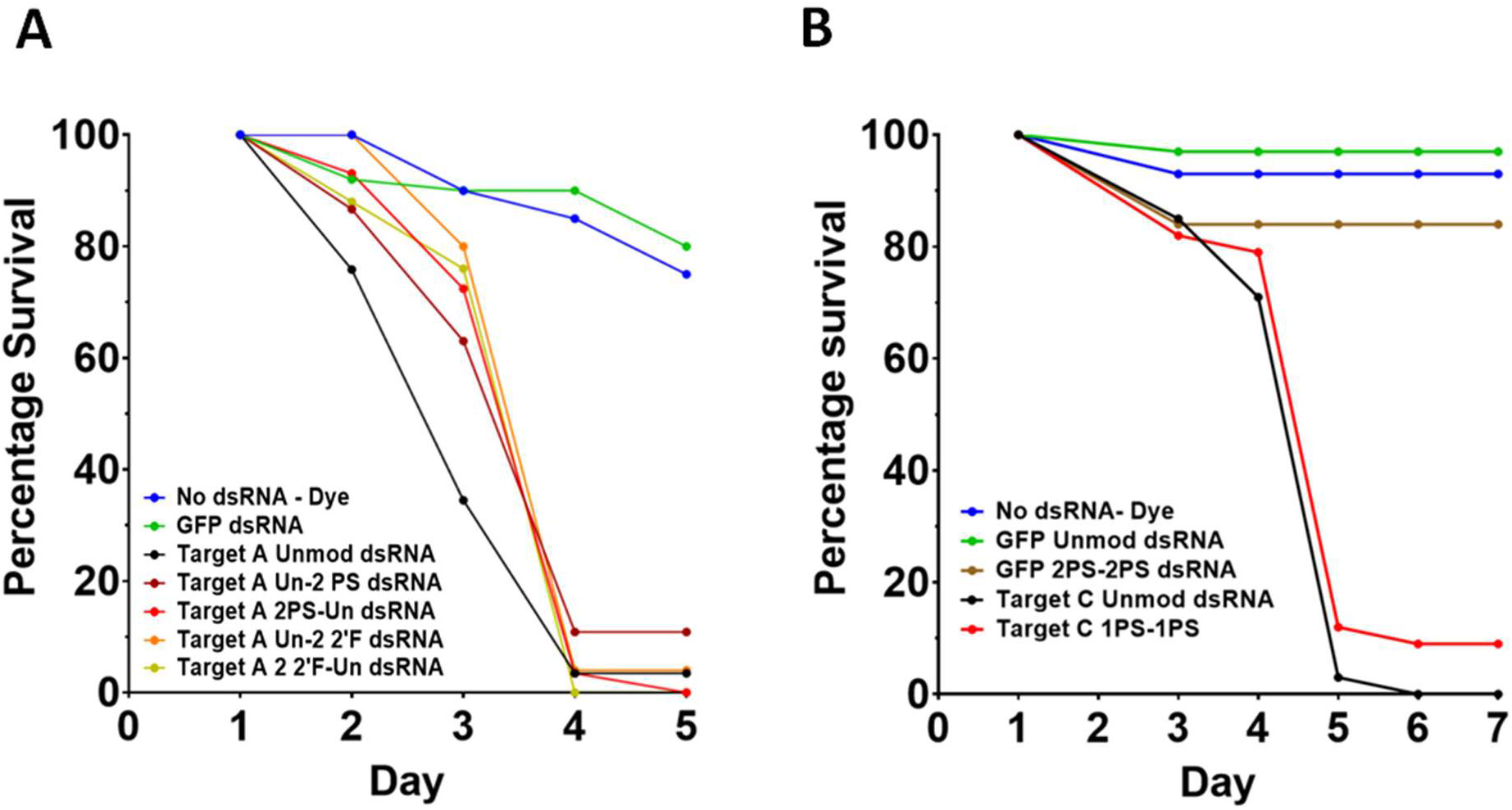
*In vivo* analysis of the effects of RNA chemical modifications of dsRNA delivered by microinjection on RNAi in stink bugs. (**A**) SGSB injection assay. Second instar (N2) insects injected on the underside of the abdomen with dsRNA solution at a concentration of 0.7 μg/ μl. Mortality measured over 5 days and normalised to day 1. Day 1 post-injection survival n – Unmod: = 29; Un-2PS: = 46; 2PS-Un: = 29; Un-2 2’F: = 25; 2’F-Un: = 25; GFP dsRNA: = 50; Dye: = 20. (**B**) SGSB injection assay. Second instar (N2) insects were injected with 10 nl of dsRNA at 1.0 μg/μl. Mortality measured over 7 days and normalised to day 1 mortality. Day 1 post-injection survival n – No dsRNA: = 29; GFP Unmod: = 37; GFP 2PS: = 37; Target C Unmod: = 37; Target C 1PS-1PS: = 39.

### Chemically modified dsRNA can induce RNAi resulting in mortality, *in vivo* in western corn rootworm (*Diabrotica virgifera virgifera*) when delivered orally in an artificial diet

Following successful induction of RNAi and insect mortality using chemically modified dsRNA in N2 stage SGSB, further insect bioassays were conducted using western corn rootworm (WCR) larvae. WCR larvae were chosen as a model organism as they are a highly relevant potential target crop pest insect for a dsRNA based insecticide, and their size allows for small controlled doses of dsRNA at a range of concentrations to be delivered orally in high throughput screening assays, to analyse RNAi efficacy of different dsRNA.

General toxicity of the chemically modified dsRNA, independent of its RNAi efficacy, was ruled out by synthesising non-targeting GFP dsRNA with PS, 2’F and HMr modifications, which were then tested in a WCR diet plate feeding bioassay. The survival results from the diet plate feeding assay after seven days, using 0.1 μg of dsRNA per well and performed in duplicate, are shown in Figure 7A. Chi-square tests demonstrate that there was no significant difference in insect mortality between insects in wells treated with chemically modified GFP, and those with unmodified GFP dsRNA or control wells with no dsRNA. In contrast, an unmodified targeting dsRNA (Target B Unmod) demonstrated high levels of insecticidal activity. The results indicate that the chemical modifications present in the dsRNA used in this experiment are not toxic to the insects and result in no significant increase in mortality above the mortality for unmodified non-targeting dsRNA. A further nontargeting control experiment was performed using a Target B scrambled dsRNA, GFP dsRNA and the Target B unmodified dsRNA as a positive control. The Target B scrambled control dsRNA showed the same low levels of insect mortality as the GFP dsRNA, whereas Target B unmodified dsRNA resulted in dose-dependent mortality as seen previously (Figure 7B and Supplementary Figure S9). This provides further confidence that mortality as a result of exposure to Target B unmodified dsRNA is due to an RNAi effect rather than a cytotoxic effect due to the base composition, and further validates the use of the non-targeting GFP dsRNA as a negative control across the different assays in this study.

**Figure 7.**
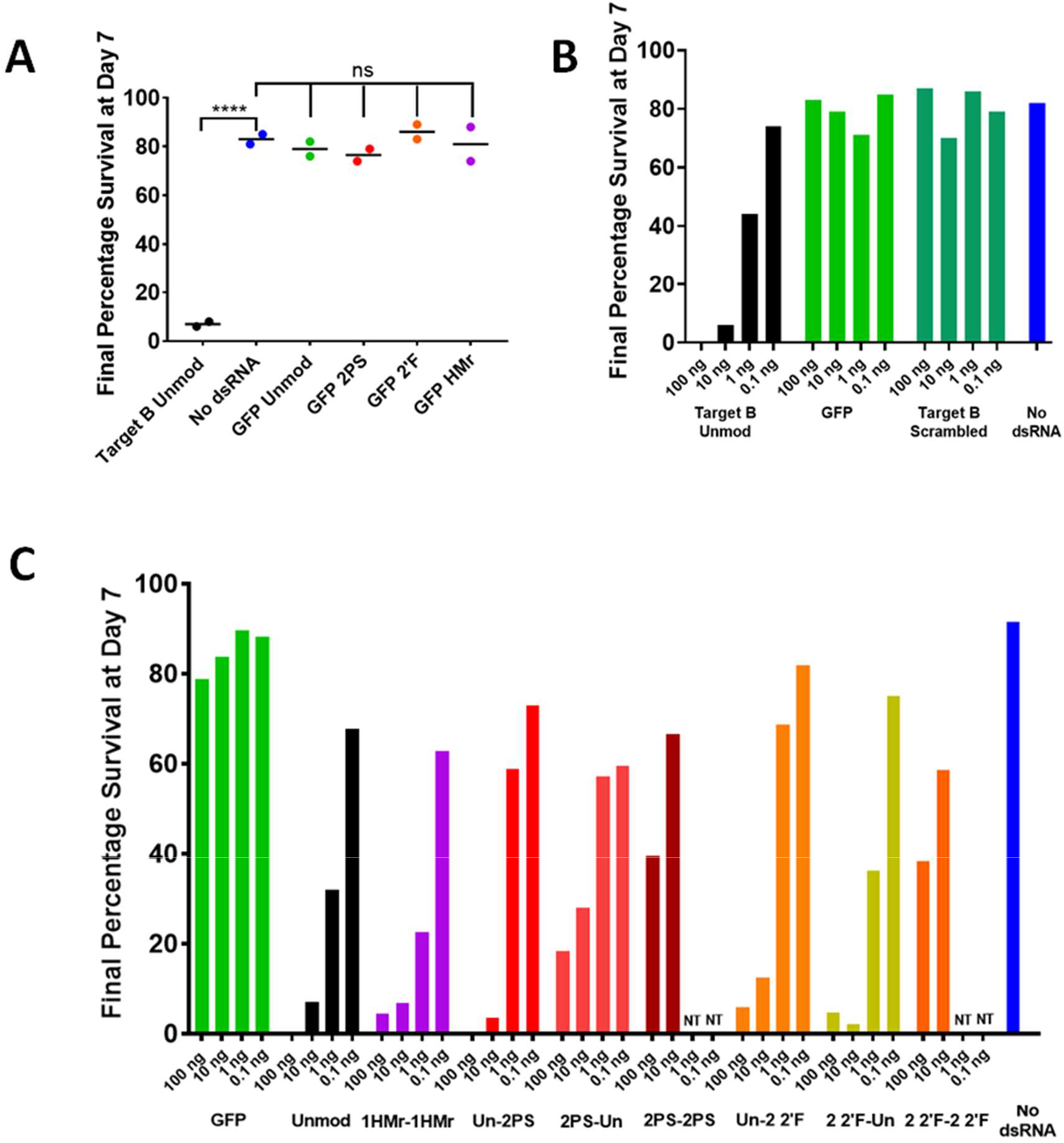
*In vivo* screening of the effects of RNA chemical modifications of dsRNA delivered in an artificial diet on RNAi in western corn rootworm. (**A**) WCR chemically modified GFP non-targeting dsRNA plate feeding assay, day 7 survival results. L1/L2 WCR larvae were fed on an artificial diet coated with 0.1 μg/well of dsRNA solution. Mortality measured over 7 days and normalised to day 1 mortality. Results of two replicate assays (N = 2) plotted, with line denoting the mean. Stars denote significance as calculated by Chi-square tests. ns = P > 0.05, **** = P ≤ 0.0001. Target B Unmod *n* = 49, 53; GFP Unmod *n* = 50, 52; GFP 2PS *n* = 50, 54; GFP 2 2’F *n* = 48, 56; GFP 1HMr *n* = 49, 46; No dsRNA *n* = 97, 98. (**B**) WCR survival feeding assay using Target B scrambled control dsRNA, day 7 survival results. L1/L2 WCR larvae were fed on an artificial diet coated with dsRNA solutions of a range of concentrations. Mortality measured over 7 days and normalised to day 1 mortality. Unmod dsRNA n = 55, 53, 52, 46, GFP dsRNA n = 51, 51, 53, 56, Scrambled dsRNA n = 48, 54, 51, 54, No dsRNA n = 114. (**C**) WCR PS, 2’F and HMr modified dsRNA plate feeding screening assay, day 7 survival results. WCR fed on an artificial diet containing dsRNA at four different concentrations (concentrations given as ng/well) in well plates. Mortality measured over 7 days and normalised to day 1. Number of insects used for each dsRNA concentration (L-R): GFP *n* = 50, Unmod *n* = 47, 48, 54, 45, 1HMr-1HMr *n* = 53, 58, 46, 41, Un-2PS *n* = 49, 48, 51, 62, 2PS-Un *n* = 51, 52, 54, 60, 2PS-2PS *n* = 48, 36, Un-2 2’F *n* = 52, 66, 38, 43, 2 2’F-Un *n* = 49, 58, 47, 48, 2 2’F-2 2’F *n* = 41, 46, No dsRNA *n* = 46.

Following optimisation of the WCR dsRNA diet plate feeding assay using unmodified dsRNA, a range of PS, 2’F and HMr chemically modified dsRNA were used in mortality assays and compared to unmodified dsRNA, with non-targeting GFP dsRNA, no dsRNA as controls. Initially, dsRNA concentrations of 100, 10, 1, and 0.1 ng per well were used, in order to ascertain which chemically modified dsRNA had similar or different activity to unmodified dsRNA. 2PS-2PS and 2 2’F-2 2’F dsRNA were only tested at the highest two concentrations, as a test assay suggested these dsRNA had greatly reduced efficacy, therefore lower concentrations would likely result in complete loss of activity (Supplementary Figures S10/11). WCR larvae were seeded onto the plates, and mortality scored and normalised (see Methods). Full mortality curves are shown in Supplementary Figure S12, and a summary of the day seven final survival results are shown in Figure 7C.

The majority of RNAi-induced mortality occurred between days 3 and 6, though our analysis was focused on the final day 7 mortality. The results for 5-hydroxymethyl modified dsRNA demonstrate that the presence of 1HMr modifications in both strands of the dsRNA did not affect RNAi efficacy as measured by insect mortality, as the final percentage survival of HMr dsRNA treated insects was similar to those of unmodified dsRNA at the two concentrations tested.

In contrast, the results for PS and 2’F dsRNA demonstrate clear differences in survival profile and final percentage survival compared to the unmodified dsRNA. Moreover, differences in RNAi efficacy as measured by insect mortality were evident depending on whether the modifications were present in the antisense strand (the intended guide strand) or sense strand (the intended passenger strand). Feeding of the dsRNA with PS modifications present only in the sense strand (Target B Un-2PS) resulted in a similar final percentage survival to unmodified dsRNA at 100 and 10 ng. dsRNA with PS modifications in the sense strand is therefore effective at inducing RNAi. Insects fed dsRNA with PS modifications present only in the antisense strand (Target B 2PS-Un) demonstrated moderate insecticidal activity at 100 and 10 ng, however, was less efficacious than unmodified and Un-2PS dsRNA. At the lower two concentrations of 1 and 0.1 ng per well, Target B Un-2PS and 2PS-Un dsRNAs showed a loss of RNAi activity compared to unmodified dsRNA, with survival of approximately 60% or above, compared to approximately 30% and 65% for 1 and 0.1 ng of unmodified dsRNA respectively.

Insects fed dsRNA with PS modifications in both strands (Target B 2PS-2PS) demonstrate a further reduction in the RNAi efficacy of the dsRNA, resulting in insect survival of approximately 40% and 65% for 100 and 10 ng of dsRNA per well respectively. PS modified dsRNA demonstrates a trend of increasing RNAi efficacy as follows: 2PS-2PS < 2PS-Un < Un-2PS < unmodified.

The results for insects fed 2’F modified dsRNA show that the overall RNAi efficacy of dsRNA with 2’F modifications in only one strand of dsRNA (Un-2 2’F, 2 2’F-Un) was similar to that of unmodified dsRNA at the highest two concentrations, though reduced slightly at lower concentrations for dsRNA with 2’F modifications in the sense strand (Un-2 2’F). In contrast, dsRNA with 2’F modifications in both strands (2 2’F-2 2’F) showed a marked reduction in RNAi efficacy similar to 2PS-2PS dsRNA, with survival of 40% and 60% respectively for 100 and 10 ng of dsRNA.

Further analysis was performed by combining data analysing 10 ng of each dsRNA from replicate experiments (Supplementary Figures S10/11) and plotted along with the mean of the two replicates (Figure 8A). The combined replicates and average confirm the previous trends in RNAi efficacy, for this assay. The results show that Un-2PS, 2 2’F-Un, and 1HMr-1HMr dsRNAs all have similar insecticidal activity to unmodified dsRNA, although the variation between replicates for Un-2PS suggests it may have a slightly reduced efficacy. 2PS-Un and Un-2 2’F have intermediate insecticidal efficacy and are statistically significantly different from their counterpart one-strand-modified dsRNAs. 2 2’F-2 2’F dsRNA has low insecticidal activity, however it is still significantly more efficacious than non-targeting GFP dsRNA, whereas the level of mortality induced by 2PS-2PS dsRNA is not significantly different from that of GFP dsRNA, suggesting 2PS-2PS dsRNA has no effective insecticidal activity at this concentration.

**Figure 8.**
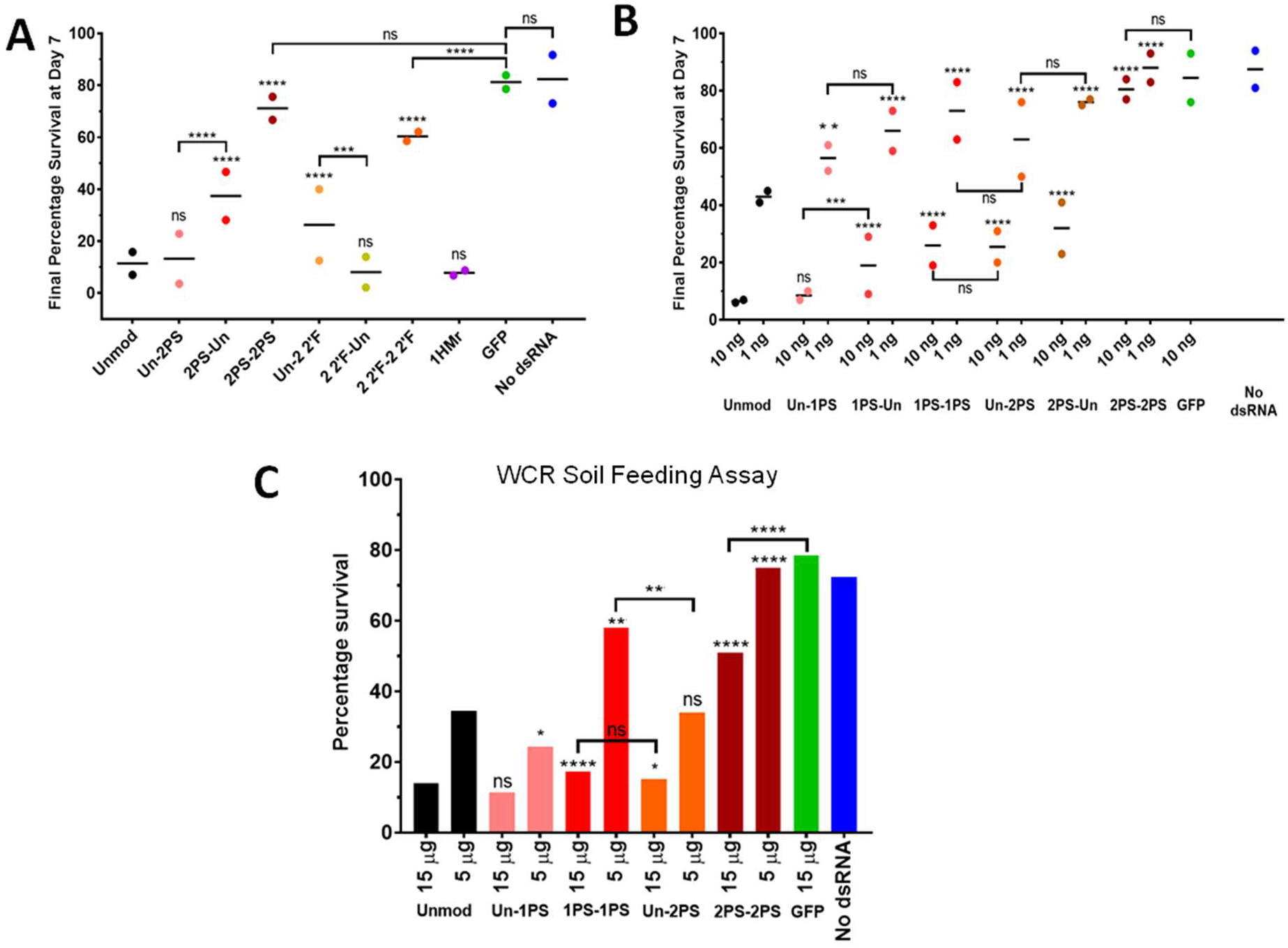
*In vivo* analysis of the effects of RNA chemical modifications of dsRNA delivered in an artificial diet and in agricultural soil on RNAi in WCR. (**A, B**) WCR chemically modified dsRNA plate feeding assays, day 7 survival results. WCR fed on an artificial diet containing dsRNA (concentrations given as ng/well) in well plates. Mortality measured over 7 days and normalised to day 1. (**A**) Combined replicates and average for 10 ng/well dsRNA from (a) and Supplementary Figure S11. Individual values are plotted along with mean indicated by a horizontal line. *N* = 2. Stars denote significance as calculated by Chi-square tests. Statistical significance symbols without brackets refer to comparison with Unmod. (**B**) Combined replicates and average for plate feeding assay with a range of 1PS and 2PS modified dsRNAs at two different concentrations. Individual values are plotted along with mean indicated by a horizontal line. *N* = 2. Number of insects used for each dsRNA concentration (L-R): Unmod Rep 1 *n* = 47, 43, Rep 2 *n* = 49, 54; Un-1PS Rep 1 *n* = 45, 52, Rep 2 *n* = 47, 49; 1PS-Un Rep 1 *n* = 48, 48, Rep 2 *n* = 46, 48; 1PS-1PS Rep 1 *n* = 48, 49, Rep 2 *n* = 50, 48; Un-2PS Rep 1 *n* = 55, 54, Rep 2 *n* = 50, 42; 2PS-Un Rep 1 *n* = 44, 50, Rep 2 *n* = 49, 48; 2PS-2PS Rep 1 *n* = 49, 48, Rep 2 *n* = 46, 45; GFP Rep 1 *n* = 47, Rep 2 *n* = 50; No dsRNA Rep 1 *n* = 96, Rep 2 *n* = 97. Stars denote significance as calculated by Chi-square tests. Statistical significance symbols without brackets refer to comparison with Unmod. (**C**) WCR 1PS and 2PS modified dsRNA soil feeding assay day 7 survival results. WCR were left on soil containing dsRNA for 1 day, then transferred to untreated diet plates and mortality measured until day 7. Number of insects used for each dsRNA concentration (L-R): No dsRNA *n* = 102; GFP dsRNA *n* = 117; Unmod *n* = 115, 119; Un-1PS *n* = 102, 116; 1PS-1PS *n* = 125, 116; Un-2PS *n* = 124, 114; 2PS-2PS *n* = 106, 121. Stars denote significance as calculated by Chi-square tests. Statistical significance symbols without brackets refer to comparison with Unmod. ns = P > 0.05, * = P ≤ 0.05, ** = P ≤ 0.01, *** = P ≤ 0.001, **** = P ≤ 0.0001

The total amount of modifications in a dsRNA with both strands modified is approximately double that of a dsRNA with only one strand modified. However, the loss in RNAi efficacy from the presence of PS or 2’F modifications in both strands is not a summation of the effects of modifications in either strand alone. This suggests that RNAi efficacy may decrease rapidly beyond a particular level of chemical modification of the dsRNA. This was investigated by focusing on dsRNA with different quantities of PS modifications (1PS and 2PS) in the sense, antisense, or both strands. For example, the total quantity of PS modifications in a dsRNA with a single nucleotide replaced by a phosphorothioate analogue throughout both strands (1PS-1PS) is likely to be approximately equal to the number of modifications in a dsRNA with two of the four nucleotides replaced by phosphorothioate analogues throughout only one strand (Un-2PS or 2PS-Un). However, the modifications are distributed very differently throughout the dsRNA and may have very different efficacies as a result.

To further analyse the effects of PS modifications in a dsRNA molecule on RNAi efficacy, a wider range of dsRNA including Un-1PS, 1PS-Un and 1PS-1PS alongside the previously studied Un-2PS, 2PS-Un and 2PS-2PS dsRNA were synthesised for use in a WCR plate feeding assay. For each dsRNA, 10 and 1 ng of dsRNA per well were used, in conjunction with GFP dsRNA and wells containing no dsRNA as controls. The assay set up, scoring, and normalisation were conducted as before.

Two replicate assays were performed and the full survival curve results are shown in Supplementary Figure S13 (1PS) and Supplementary Figure S14 (2PS), and final day seven percentage survival summarised in Figure 8B. Variation was observed at some time points, particularly around day 3, however, the replicates show good agreement on final day 7 percentage survival (Figure 8B).

The results show that for both 1PS and 2PS dsRNA, dsRNA with PS modifications in the sense strand (Un-1PS, Un-2PS) showed slightly increased RNAi activity compared to their respective counterparts with PS modifications in the antisense strand (1PS-Un, 2PS-Un), though the difference was only statistically significant for Un-1PS and 1PS-Un dsRNA at 10 ng. However, only dsRNA with 1PS modifications in the sense strand had similar RNAi efficacy compared to unmodified dsRNA; all other PS modified dsRNAs showed reduced RNAi efficacy compared to unmodified dsRNA. Furthermore, increasing the number of PS modifications in the sense strand resulted in reduced RNAi activity.

2PS-2PS dsRNA was markedly less active than 2PS-Un dsRNA or any other targeting dsRNA, again demonstrating no significant difference in insecticidal activity compared to GFP control dsRNA. These results demonstrate a progressive decrease in RNAi activity due to the increasing number of PS modifications. The previous observations for 2PS modified dsRNA regarding the link between RNAi efficacy and which strand of the dsRNA contains PS modifications were also demonstrated to be true for 1PS modified dsRNA. It was noted that the rank order of RNAi efficacy for the 2PS dsRNA was the same as in the previous screens. However, the efficacies of the 2PS dsRNA were lower compared to some previous experiments.

### Unmodified and phosphorothioate dsRNAs demonstrate insecticidal activity when applied to biologically active soil containing nucleases

The results of the WCR feeding screens conducted in diet plates, showed that although a number of phosphorothioate modified dsRNAs had similar RNAi efficacy compared to unmodified dsRNA, none demonstrated improved RNAi efficacy. Therefore, further experiments were performed to determine if the potential benefits in improved resistance of the PS modified dsRNA to both insect and soil nucleases (Figures 2/3) impacted on overall RNAi efficacy. As WCR larvae feed on the roots of corn plants, their environment is the surrounding soil. Soil contains a variety of live and decaying microorganisms, arthropods, and plant matter, all of which are sources of extra-cellular nucleases capable of degrading insecticidal dsRNA prior to it being ingested by the target insect. Protection against the activity of these nucleases by phosphorothioate modifications as seen in Figures 2/3, could therefore ensure insecticidal phosphorothioate dsRNA would remain intact and active in the soil for a greater length of time compared to unmodified dsRNA. 2’F dsRNA also demonstrated increased soil nuclease resistance, however demonstrated variable RNAi efficacy between *in vitro* and *in vivo* screening assays and so was not tested.

In order to test the hypothesis that phosphorothioate dsRNA would maintain their insecticidal activity in soil for longer than unmodified dsRNA, unmodified and various phosphorothioate dsRNA were used in a soil feeding assay. This assay involved the application of dsRNA to the same defined, biologically active soil used in the soil nuclease assay. This contains naturally occurring live microorganisms and active nucleases, within which WCR larvae were then placed for around 24 hours. The larvae were then transferred to diet plates, and their mortality scored and normalised as for the diet plate feeding assay. The diet plate feeding assay had conclusively shown that phosphorothioate modifications in the antisense strand reduced the RNAi activity compared to unmodified dsRNA, and thus insecticidal activity of dsRNA, therefore were not used in this study. Modifications in both strands also reduced RNAi activity, however data from the stink bug saliva nuclease stability assay indicated that modifications in both strands increased nuclease stability compared to modifications in a single strand, or unmodified dsRNA (Supplementary Figure S3). dsRNA with phosphorothioate modifications in both strands were therefore also used, as the increased stability of the dsRNA in the soil might be a greater contributing factor to overall insecticidal activity compared to any reduction of RNAi activity due to the chemical modifications.

The selected unmodified, 1PS and 2PS dsRNA were analysed across a concentration range 0.6-15 μg of dsRNA per soil sample. dsRNA was added to two soil samples for each concentration of each dsRNA. The results are shown in Figure 8C and Supplementary Figures S15/S16. Un-1PS and Un-2PS dsRNA demonstrated similar insecticidal activities to unmodified dsRNA at 15 and 5 μg, as did 1PS-1PS dsRNA at the highest concentration of 15 μg. At 5 μg Un-1PS demonstrated efficacy 10% above that of unmodified dsRNA, although the difference was not significant. At 15 μg 2PS-2PS dsRNA also has low to moderate insecticidal activity, significantly above that of GFP control dsRNA. Also of note was the greater insecticidal activity demonstrated by Un-2PS dsRNA compared to 1PS-1PS dsRNA at 5 μg.

## DISCUSSION

We synthesised a range of chemically modified long dsRNA substrates to study the effects of these chemical modifications on nuclease resistance and RNAi efficacy both *in vitro* and *in vivo* in insects. This was with a view to using chemical modifications to improve the efficacy of dsRNA insect control. The results obtained demonstrate that long chemically modified dsRNA which target insect mRNAs can be successfully synthesised *in vitro*, and a number of the chemically modified dsRNA resulted in improved resistance against insect and environmental nucleases. We demonstrated – to our knowledge, for the first time – that long chemically modified dsRNA are effective triggers for RNAi in a *Drosophila* (Diptera) insect cell line (Kc167 cells), live southern green stink bug nymphs (Hemiptera), and live western corn rootworm (WCR) larvae (Coleoptera). The results also demonstrate that in the quantitative cell-based assay, several of the chemically modified dsRNA including 1PS, 2PS and 1 2’F modified dsRNA, demonstrated improved RNAi efficacy compared to unmodified dsRNA.

The overall RNAi efficacy of a chemically modified dsRNA depends on a complex combination of how the chemical modifications affect environmental and insect nuclease degradation; cellular dsRNA uptake and trafficking; Dicer binding and cleavage to esiRNAs; RISC binding, duplex unwinding, and strand selection; and target mRNA binding and cleavage. The results presented in this study, provide further mechanistic insight into the effects of a number of chemical modifications on dsRNA used for insect control and the potential advantages for future applications.

Improving the stability of dsRNA to environmental nucleases, including nucleases present in the soil, insect saliva, gut secretions, and hemolymph are potential strategies that could be utilised to improve the RNAi efficacy of RNA based products. Nuclease stability studies using stink bug saliva nucleases and soil nucleases showed that phosphorothioate dsRNA had increased resistance when compared to unmodified dsRNA. In addition, 2’-fluoro modified dsRNA demonstrated increased resistance against degradation by soil nucleases. These results are consistent with previous studies which showed the increased half-life and resistance of phosphorothioate and 2’-fluoro siRNAs and antisense oligonucleotides to degradation by mammalian nucleases (39–41, 48). These results for the first time demonstrate the ability to improve the stability of dsRNA towards insect and/or agricultural environmental nucleases using phosphorothioate and 2’-fluoro modifications. However, none of the chemically modified dsRNAs used in this study demonstrated increased resistance towards CPB gut nucleases, or environmental degradation as a result of UV radiation exposure.

RNA chemical modifications that increase nuclease stability, may reduce the RNAi efficacy of a dsRNA by hindering key steps in the RNAi pathway itself including inhibiting Dicer processing of long dsRNA into esiRNAs. Indeed, the results of our in vitro Dicer/RNase III cleavage assays demonstrate that the ability of these enzymes, to process dsRNA to esiRNAs can be affected by certain chemical modifications, and variation in the ability of Dicer to cleave some chemically modified dsRNA may explain differences in RNAi efficacy seen between insect species, or between live insect and *in vitro* insect cell RNAi assays. However, previous studies using small chemically modified dsRNA duplexes showed Dicer processing is not required for incorporation into RISC and gene silencing (49). Therefore, it may be that some fragments of the long dsRNA that are larger than the typical 21-25 bp size of canonical siRNAs can still be incorporated into functional silencing complexes.

The analysis of the effect of chemical modifications of long dsRNA on bacterial RNase III and Dicer processing *in vitro*, showed that bacterial RNase III was able to cleave dsRNA containing PS, 2’F and HMr modifications into siRNAs. These results are consistent with previous observations demonstrating that bacterial RNase III cleavage specificity is not altered by phosphorothioate bonds (50). Results of *Giardia* Dicer processing of long dsRNA containing PS, 2’F and HMr modifications demonstrated that the presence of these modifications reduced the ability to cleave the long dsRNA into esiRNAs. These results are consistent with previous studies using small dsRNA duplexes (25 bp) containing minimal 2’-O-Me modifications inhibited the ability of human recombinant Dicer to process the dsRNA to 21 bp siRNA products (49). In addition, using 27 bp dsRNA the use of 2’-methoxyribonucleotide substitutions at the cleavage sites also prevented cleavage *in vitro* human dicer cleavage assays (51). However, *Giardia* Dicer has been shown to cleave dsRNA with 2’-fluoro modifications (44). The observed differences in processing long chemically modified dsRNA substrates is potentially reflected by differences between bacterial RNase III and Dicer. RNase III binds dsRNA with a dsRNA binding domain (dsRBD), whereas *Giardia* Dicer binds 2 nt ssRNA overhangs using only a PAZ domain (52, 53) The PAZ domain of *Giardia* Dicer specifically binds the 3’ two-base overhang at the end of a dsRNA substrate (54) and previous studies have demonstrated that the 3’ terminus is critical in the binding interaction of the enzyme with the substrate and that this process is sensitive to base composition (51). Bacterial RNase III has a single RNase III domain and function as homodimers (53, 55, 56) whereas Human Dicer and Drosha have two RNase III domains in the same molecule and function as monomers (54, 57). Recently it was shown that bacterial RNase III cleaves long dsRNA into a set of small RNAs with distinct patterns of length distribution (58). Additionally, Dicer nucleases with similar domain architectures such as human Dicer and *Drosophila* Dicer-2 (see Supplementary Figure S5) will have significant differences in domain size and amino acid composition that may result in variations in their ability to bind and cleave chemically modified dsRNA.

The RNAi efficacy of the chemically modified long dsRNA was studied both *in vitro* in *Drosophila* insect cells in and *in vivo* in live insect feeding assays. The results of the quantitative dual luciferase assays showed that long dsRNA with chemical modifications in both the antisense and sense strands are potent RNAi inducers in *Drosophila* cells (see Figure 5). Moreover, the results demonstrate that dsRNA containing 2 phosphorothioate modifications in both strands (2PS-2PS) results in improved RNAi efficacy compared to unmodified dsRNA (IC50: 2.8 ng compared 20.6 ng). Further studies to analyse RNAi efficacy based on insect mortality using chemically modified dsRNA in N2 stage SGSB showed that the chemically modified dsRNA used were all effective, resulting in similar insect mortality after 7 days compared to the unmodified dsRNA (see Figure 6). *In vivo* experiments using WCR larvae where RNAi efficacy was measured by insect mortality, also showed that the RNAi efficacy for both the 1PS dsRNA or 2 2’F dsRNA with modifications in only one strand was similar to unmodified dsRNA. However, further increasing the amount of the chemical modification present (2PS dsRNA and 2 2’F dsRNA with both strands modified) resulted in a significant reduction in RNAi efficacy, and hence almost total loss of observable insect mortality. These findings were unexpected given the improved RNAi efficacy observed in *Drosophila* Kc167 cells and the increased resistance observed for 2PS dsRNA to stink bug saliva nucleases.

There is clearly a difference in how the heavily modified phosphorothioate dsRNA (2PS-2PS) affect cellular uptake, Dicer-2 processing, and/or RISC assembly between insect orders as efficient RNAi was achieved using 2PS-2PS (2PS) dsRNA in *Drosophila* cells (from the order Diptera) in contrast to the RNAi and low insecticidal activity in WCR larvae (from the order Coleoptera). Variations observed for the same type of chemically modified dsRNA, against different nucleases and in RNAi efficacy between different insect species or orders, demonstrate that there is not a universal mechanism in which these chemical modifications modulate overall RNAi efficacy in all situations or species. Rather, use of RNA chemical modifications to improve the efficacy of dsRNA insecticides is highly specific to the application environment (e.g. leaf or soil) and the target species, in much the same way that delivery and uptake of unmodified dsRNA varies between species, and delivery methods (21, 22). Where dsRNA insecticides are in the process of being developed commercially, a great deal of time goes into bioinformatics, including sequence design and bioassay testing of effective target mRNAs and identification of mRNA target regions that allow the dsRNA insecticide to be highly selective. In addition, formulations are developed that stabilise dsRNA or aid in their uptake by the target insect. It is therefore conceivable that tailoring the type, and strand distribution of chemical modifications in a dsRNA insecticide, could be an additional step in the developmental process of dsRNA based plant protection

Results in this study on the effects of the chemical modifications in either the antisense or sense strands also provide further mechanistic insight. Where the RNAi efficacies of dsRNAs with chemical modifications in either of the two strands but not both strands are similar to each other but different from unmodified dsRNA, this suggests Dicer processing or uptake may be affected by the modifications, but RISC loading is unaffected. Conversely, where one of the dsRNAs with chemical modifications in one strand is more efficacious than the other, this suggests the modification may primarily affect strand selection or RISC loading, though differences in efficacy from unmodified dsRNA may include components due to uptake or Dicer processing as well.

For example, the results from the WCR feeding assay showed a greater reduction in RNAi efficacy when the phosphorothioate modifications are present in the antisense strand. This is clearly observed in the WCR feeding assay for 2PS-Un dsRNA compared to Un-2PS and (see Figure 7). These results suggest that in the live insects, chemical modifications in the antisense strand may affect RISC assembly, discarding of the sense (passenger) strand, and subsequent use of the antisense strand to bind the mRNA target and resulting mRNA cleavage. These results are consistent with previous studies where mammalian RNAi machinery has demonstrated preferences for chemical modifications in one strand of an siRNA duplex. siRNAs containing chemical modifications in the sense strand have been shown to demonstrate greater RNAi efficacy than those with chemical modifications in the antisense strand (59, 60).

In contrast, dsRNA with 2’F modifications in the sense strand showed reduced RNAi efficacy compared to that of unmodified dsRNA in WCR. This suggests that selection, binding, and discarding of the sense strand (the intended passenger strand) during loading of the antisense strand (the intended guide strand) into the RISC may be affected by 2’F modifications. Increasing the level of chemical modifications in both strands (2PS-2PS and 2’F-2’F) resulted in a significant reduction in RNAi efficacy as measured by insect mortality. This result, in conjunction with the results of the in vitro Dicer/RNase III assays, suggests that this may inhibit Dicer-2 processing of the long dsRNA which may also contribute to reduced RNAi efficacy of the 2PS-2PS dsRNA compared to the 2PS-Un dsRNA observed in WCR. The further reduction in RNAi efficacy for 2PS-2PS and 2’F-2’F dsRNA may also be a result of increased issues with RISC loading, or increased difficulty with duplex unwinding due to changes in thermal stability. Further investigation is required to determine if one or several of these are major factors in overall RNAi efficacy of a chemically modified dsRNA in insects.

These results demonstrate the potential for the application of chemically modified long dsRNA based insect control. Further studies analysing the effects of a wider range of chemical modifications (e.g. boranophosphates, 2’-O-methyl, phosphorodithioate and LNA) may generate further improvements in the resistance towards insect and environmental nucleases and demonstrate improved RNAi efficacy *in vivo*. Chemical modification of long dsRNA provides a number of potential advantages, including increased stability and specificity for the development of long dsRNA actives for applications in insect management strategies. Further work is required in order to determine how chemically modified dsRNA result in different RNAi efficacies in different insect species, or between cell-based and live insect systems, and which steps of the RNAi pathway in insects are responsible for the differences observed. These additional insights could help in the future development of chemically modified dsRNA insect control.

## EXPERIMENTAL PROCEDURES

### Synthesis of dsRNA

DNA templates for in vitro transcription of dsRNA were produced by polymerase chain reaction (PCR). DNA templates were generated with either a single T7 RNA polymerase promoter to synthesise ssRNA templates, or two T7 RNA polymerase promoters in opposing directions to generate dsRNA. PCR was performed using 12.5 μl KAPA2G Fast PCR mastermix (KAPA Biosystems), containing reaction buffer, MgCl2, dNTP mix and DNA polymerase; 1 μl of initial DNA template (approximately 10-50 ng); 1.25 μl each of 10 μM forward and reverse primers (IDT) (see Supplementary Table 1). PCR reactions used the following conditions: 95 °C for 3 mins; 30 cycles of: 95 °C for 15 secs, 60 °C for 15 secs, 72 °C for 3 secs; 72 °C for 1 min. Final PCR DNA products were purified using a Qiagen QIAquick PCR Purification Kit following the kit protocol, and used as the DNA templates for subsequent IVT reactions.

IVT reactions were performed using the Ambion MEGAscript T7 IVT kit (ThermoFisher). For unmodified ss/dsRNA: 2 μl of each the 75 mM NTPs, 2 μl of 10 X reaction buffer, 0.1-0.3 μg DNA template, and 2 μl MEGAscript T7 polymerase, made up to 20 μl with nuclease-free water.

For phosphorothioate (PS) ss/dsRNA: IVT reactions performed as described for unmodified with 2 μl of the appropriate unmodified 75 mM NTP replaced by 5 μl of 10 mM Sp α-thiophosphate NTP (Biolog) or 2 μl 100 mM Rp/Sp α-thiophosphate NTP (TriLink).

For 5-Hydroxymethyl ss/dsRNA: IVT reactions performed as described for unmodified RNA, however 2 μl of the appropriate canonical NTP (75 mM) was replaced by 2 μl of 100 mM 5-Hydroxymethyl-5’-triphosphate (e.g. 5-Hydroxymethyl-CTP) (TriLink).

2’-Fluoro ssRNA was produced by IVT using the Durascribe T7 IVT kit (Epicentre). Reactions consisted of 0.2-1 μg of purified DNA template per 20 μl reaction, with 2.0 μl of each 50 mM NTP or 2’F-NTP, 2 μl 10 X reaction buffer, 2 μl 100 mM DTT and 2 μl Durascribe T7 polymerase, made up to 20 μl with nuclease-free water.

All IVT reactions were incubated at 37 °C for 4-16 hours, prior to removal of DNA template by addition of 1 μl DNase (Ambion MEGAscript kit, Durascribe kit) per 20 μl IVT reaction mixture, and incubation at 37 °C for 20 mins. ss/dsRNA was purified by solid phase extraction (SPE) as previously described (61). Quantification was performed using a Nanodrop 2000 UV visible spectrophotometer (Thermo Fisher Scientific). Extinction coefficients ((μg/ml)^-1^ cm^-1^): dsDNA – 0.020, ssRNA – 0.025, dsRNA – 0.021; Concentration (μg/ml) if A260 value = 1: dsDNA – 50, ssRNA – 40, dsRNA – 46.52 (62). dsRNA annealing was performed using equal quantities of ssRNAs (1-300 μg) in 1 X PBS. The mixture was heated to 85 °C for 2-4 mins and then allowed to cool to room temperature.

### Ion pair reverse phase high performance liquid chromatography (IP RP HPLC) analysis of RNA

Samples were analysed by IP RP HPLC on a passivated Agilent 1100 series HPLC using a Proswift RP-1S Monolith column (50 mm x 4.6 mm I.D. ThermoFisher). Chromatograms were generated using UV detection at a wavelength of 260 nm. The chromatographic analysis was performed using the following conditions: Buffer A - 0.1 M triethylammonium acetate (TEAA) pH 7.0 (Fluka), 0.1 % acetonitrile (ACN) (ThermoFisher); Buffer B - 0.1 M TEAA pH 7.0, 25 % ACN. RNA/DNA was analysed using the following gradient. Gradient starting at 20% buffer B to 30% in 1 minute, followed by a linear extension to 70% buffer B over 11.5 minutes, then extended to 100% buffer B over 1 minute, held at 100% buffer B for 2 minutes, reduced to 20% in 0.1 minutes and held at 20% for 4.5 minutes at a flow rate of 0.75 ml/min at 50 or 75 °C, with temperature controlled by an external column oven (Transgenomic).

### Stink bug saliva nuclease degradation assay

Saliva was collected in feeding sachets produced by vacuum pumping Parafilm over a 96 well plate. The resulting indentations over wells were loaded with 25 μl Sf-900 insect cell culture media (Gibco), sachets sealed, and placed over 96 well mesh bottom plates containing one N2 SGSB nymph per well. The remaining saliva-containing Sf9 media was extracted by syringe after 3 days of the insects feeding, pooled, and stored at −20°C until required.

The initial saliva-containing Sf9 media was successively diluted 1:3 in MilliQ water across rows of a 96 well plate, down to 1:729, plus a water only control row for each dsRNA tested. Two technical replicates were set up for each dsRNA, with one whole set of replicates in each of two separate plates. 2.6 μg of dsRNA in 20 μl of nuclease-free water was loaded per well and the plates sealed. Plates were incubated at room temperature on a shaker plate. 20 μl samples were collected for each combination of dsRNA and saliva-media dilution at 2, 4 and 6 hours and after overnight incubation (approximately 16 hours). Collected samples were dispensed into fresh 96 well plates containing gel loading dye and stored at −20 °C until thawed for gel electrophoresis analysis.

### Agricultural soil supernatant nuclease degradation assay

Soil supernatant was prepared by mixing 0.1 g of agricultural soil with 200 μl of nuclease-free water, vortexing for 2 minutes, followed by centrifugation at 13,000 rpm for 1 minute, and collection of the supernatant. The pH of the soil in water and 0.01 M CaCl2 was determined as 7.6 and 7.4 respectively, giving confidence that any dsRNA degradation seen was due to nuclease activity in the soil and not to alkaline hydrolysis of the RNA.

Reaction solutions were made up containing 1 μg of dsRNA in 2 μl of water, 3 μl of soil supernatant for a total of 5 μl per time point (25 μl total) and reactions incubated at 37 °C. 5 μl samples were removed at each time point (0 hrs, 4 hrs, 23 hrs, 28 hrs and 48 hrs), mixed with formamide loading dye, and frozen at −20 °C prior to agarose gel electrophoresis analysis. Reactions were performed in triplicate, with three soil supernatants made from different samples of the same type of soil.

### Colorado potato beetle gut secretion nuclease degradation assay

CPB gut secretions were collected from L4 larvae by agitating their mouths with a glass capillary until gut contents was expelled, which was collected in a microcentrifuge tube incubated on ice, and frozen at −20 °C until use. Gut secretion dilutions were created by diluting in nuclease-free water. For the dilution assay, reaction solutions were made up for each dilution of gut secretion containing 1 μg of dsRNA in 2 μl of water, and 3 μl of gut secretion dilution. Reactions were incubated at 37 °C for 30 mins, then formamide loading dye was added, and samples frozen at −20 °C until gel analysis. For the time course assay, reaction solutions were made up containing 1 μg of dsRNA in 2 μl of water per time point, and 3 μl of 1/1,000 gut secretion dilution per time point. Reactions were incubated at 37 °C and 5 μl samples were removed at each time point (15 min, 30 min, 1hr, 2hr), mixed with formamide loading dye, and frozen at −20 °C until gel analysis.

### UV exposure degradation assay

Reaction solutions were made up containing 1 μg of dsRNA in 10 μl of nuclease-free water per timepoint in UV-permeable microcuvettes sealed with caps to prevent evaporation. Microcuvettes were exposed to 254 nm UV radiation in a CL-1000 100 μJ/cm^2^ UV crosslinker (UVP), and 10 μl samples removed at time points of 30 min, 1 hr, 2hr and 3 hr, formamide loading dye was added, and samples frozen at −20 °C until gel analysis.

### RNase A nuclease degradation assay

1U RNase A (Thermo Fisher) was incubated with 1 μg dsRNA in 0.5 M NaCl (10 μl total volume) for 20 mins at 37 °C followed by addition of formamide loading dye and immediate gel electrophoresis analysis.

### *In vitro* Dicer/RNase III processing assays

#### RNase III assay

1 U (0.5 μl) RNaseIII (NEB Short Cut RNase III), was combined with 1 μl of 10X reaction buffer, 1 μl of 10X MnCl2, 1 μg of dsRNA and reactions made up to 10 μl with nuclease free water. Reactions were incubated at 37 °C for 20 mins.

#### *Giardia* Dicer assay

1 U (1 μl) *Giardia intestinallis* PowerCut Dicer (ThermoFisher), was combined with 1 μl 5X reaction buffer, 1 μg of dsRNA and reactions made up to 5 μl with nuclease free water. Reactions were incubated at 37 °C for 16 hours.

### Gel electrophoresis analysis

Frozen samples from saliva nuclease degradation assays were analysed on 1% (w/v) agarose gels stained with GelRed (Biotium) and visualised on a BioRAD Gel Doc EZ Imager with a UV transilluminator. Frozen samples from all other nuclease/degradation/Dicer processing assays were analysed on 1.2% (w/v) agarose gels, and gels stained using ethidium bromide. Quantification of gel bands was performed with Fiji (ImageJ) image analysis software using the in-built gel band quantification tool. Results were given as ‘relative dsRNA stability index’. Relative dsRNA stability index = (Band intensity of dsRNA incubated with nuclease or exposed to UV)/(Band intensity of dsRNA incubated with water or sample from time point 0).

### Insect cell culture

*Drosophila* Kc167 cells were cultured in Hyclone CCM3 Insect Media (GE Life Sciences) containing 1% Pen-Strep (Lonza). Cells were thawed in medium supplemented with 10% FBS (Sigma Life Sciences), then once established, cultured in serum-free CCM3 medium with Pen-Strep for amplification and in assays. Cells were cultured in T75 flasks (Thermo Scientific) and passaged as required based on confluency.

### Dual luciferase assays

*Drosophila* Kc167 cells from a stock were seeded at around 60-70 % confluence in T75 flasks the day prior to transfection. A Qiagen Effectene Transfection Reagent kit was used as per the kit protocol, scaled to treat a T75 flask as equivalent to a 75 mm dish, with each flask receiving: 225 μl of EC buffer, 12 μl of enhancer, 45 μl of Effectene, and a total of 1.5 μg of plasmid DNA: 375 ng *Renilla* luciferase plasmid, 750 ng 6×2xDraf firefly luciferase plasmid, 225 ng Upd plasmid, 150 ng pAc5.1.

Buffer, enhancer and pDNA were combined and incubated at room temperature for 5 minutes, followed by addition of Effectene and incubated at room temperature for 8 minutes. This mixture was then made up to 1.5 ml with fresh cell culture medium without Pen-Strep. T75 flask media was replaced with 7 ml fresh medium (without Pen-Strep) and the 1.5 ml reaction solution, swirled gently to ensure mixing and incubated at 25 °C overnight.

dsRNAs were dispensed into 96 well plates, with dsRNA diluted in 20 μl of nuclease-free water, and 6 technical replicate wells per combination of dsRNA and concentration. Edge wells were filled with water and left unused in order to guard against plate edge effects.

The day after transfection the transfection medium was aspirated from the T75 flasks. Cells were resuspended in enough fresh serum-free CCM3 medium with Pen-Strep to provide 40,000 cells per well of a 96-well plate, in 100 μl. Cells were dispensed into the dsRNA-containing plates, sealed, centrifuged for 1 min at 2,000 rpm to encourage adherence of cells to well bottoms, then incubated at 25 °C for 4 days to allow RNAi knockdown of the target luciferase protein to occur.

The dual luciferase assay was performed using the Dual-Luciferase Reporter Assay System kit (Promega). 1X passive lysis buffer (PLB) was prepared from 5X concentrate. The Luciferase Assay Reagent II or LARII (FL reagent), and the Stop&Glo Reagent (RL reagent) were mixed with the buffers provided in the kit and then the solutions diluted 1:2 in MilliQ water. Plates were centrifuged for 2 mins at 2,000 rpm to ensure adherence of loose cells, then the medium was aspirated, leaving 10 μl of residual medium per well. 20 μl 1X PLB added per well and plates incubated at room temperature for 15 minutes. 100 μl of LARII dilution was added to each well and the fluorescence of each well (FL values) was read using a Varioskan Flash plate reader (Thermo Scientific) with a pre-reading 10 sec shake step. 100 μl of Stop&Glo dilution was then added to each well, and the plate re-read with the filter applied (RL values).

Unpaired T-test analysis of luciferase assay data was performed using GraphPad Prism software. Dose curves and IC50 values were generated by non-linear regression analysis using a dose-response inhibition variable slope curve, also using Graphpad Prism software.

### Insect culture

SGSB were reared on runner beans at 26 °C with 50% relative humidity on a light:dark regimen of 16 hrs:8 hrs. WCR were reared in trays containing corn plants at 26 °C with 65% relative humidity on a light:dark regimen of 16 hrs:8 hrs.

### Stink bug injection assays

N2 nymph stage SGSB were fixed to microscope slides with their undersides exposed using double sided tape, injected in the abdomen, and then liberated again using cooking oil. Insects were transferred to a sealed dish containing a single runner bean, with one dish per condition. The injection day was designated day 0; on day 1 all deceased insects were removed and the initial scoring done. It was assumed all mortality in the first day was the result of damage incurred during injection and not from RNAi. Mortality was scored over subsequent days and recorded.

The first injection assay (Figure 5A) used an Eppendorf EDOS 5222 injector and solutions of dsRNA at concentrations of 700 ng/ul. With this system the injection volume was variable depending on insect size. The second injection assay (Figure 5B) used a Drummond Nanoject III Programmable Nanoliter Injector to deliver a fixed dose of 10 nl of a 1 μg/μl dsRNA solution per insect.

### Western corn rootworm diet plate feeding assay

WCR larvae were fed on artificial diet for the duration of the assay, with 500 μl of diet set in the bottom of each well of a 48 well plate. Aqueous dsRNA solutions prepared using MilliQ water and purified and annealed dsRNAs in 1X PBS, were applied to the diet surface and plates dried in a lamina flow hood. Approximately two larvae were seeded per well, and the plate sealed, with air holes in the film to allow air exchange with the wells. Initial mortality was scored at the end of day zero. Mortality was scored each subsequent day for seven days and normalised to the survival of the first day in order to negate most of the non-RNAi mortality associated with mishandling of rootworms during plate seeding. Each experiment included control plates containing no dsRNA and GFP dsRNA. Further individual control experiments included plates containing chemically modified GFP dsRNA and a scrambled dsRNA. Half a plate was treated with each combination of dsRNA type and dsRNA concentration, resulting in approximately 48 larvae in 24 wells per condition.

WCR Artificial Diet For 1.2 L: 19.3 g of agar was dissolved in 1.2 L of autoclaved MilliQ water. 36.6 g of wheat germ, 43 g of casein and 12.3 g of Wesson salt mix sieved into agar. Corn leaf powder ground for 10 minutes in a tissuelyser (Qiagen) on maximum frequency and 8.4 g sieved into diet mix. 43 g of sucrose, 18.3 g of alpha cellulose, 2 g of vanderzant modification vitamin mixture, 1.33 g of nipagin (methylparaben preservative) and 0.83 g of sorbic acid added to the diet mix. The following were weighed using an analytical balance and added to the diet: 83 mg of cholesterol, 170 mg Aureomycin (chlortetracycline), 170 mg rifampicin, 170 mg chloramphenicol, and 67 mg nystatin. Finally, 330 μl linseed oil, and 6.6 ml of 10% (w/v) KOH were added.

### Western corn rootworm soil feeding assay

For each dsRNA, two wells of a 48 well plate were dedicated. Plates were set up in duplicate for each dsRNA, to give two time points, with one set of plates having corn rootworm larvae applied on day 0 (week 0), another set of plates had larvae applied on day 7 (week 1). Each plate had two wells each containing a base layer of 300 μl of agar, with 370 mg of Stein soil (live defined soil comprising a third each clay, silt, and sand) on top to which was applied 50 μl of dsRNA solution containing the appropriate dose of dsRNA (15, 5, 1.7 and 0.6 μg per well). Control plates with 15 μg of GFP dsRNA, and a water only dsRNA-free negative control were also set up.

130 L1 larvae were applied to each of the two wells for a total of 260 insects per combination of dsRNA, concentration, and time point. The week 1 plates were incubated in WCR rearing conditions from day 0 onwards along with the day 0 plates. After the application of insects to each plate at each of the time points, insects were left in the soil for 24 hours, then extracted from the soil and live larvae transferred into 48 well diet plates as used for the corn rootworm diet plate feeding assay with 2 to 4 larvae dispensed into each well. After each day where insects were transferred (days 1 and 8) mortality was scored over subsequent days for seven days. Survival data for each time point was normalised to the number of surviving larvae upon transfer from soil to diet plates (days 1 and 8).

### Western corn rootworm diet plate and soil feeding assays data analysis

Statistical significance was assessed using a Chi-square test of independence. Tests were performed using percentages to avoid false significance where there was a large difference in total (live + dead) n numbers between the two groups being compared, though corroborated by tests using raw n numbers of alive/dead insects. Where available, the total n numbers and the average percentage survival of two replicates were used.

## SUPPORTING INFORMATION

This article contains supporting information.

