## Supplementary for "Chemically modified dsRNA induces RNAi effects in insects *in vitro* and *in vivo*: A potential new tool for improving RNA-based plant protection"

**TABLE OF CONTENTS**

**Table S1.** Primers used in the PCR synthesis of DNA templates for IVT reactions. T7 polymerase binding sites are marked in bold.

**Figure S1.** Analysis of dsRNA synthesised by *in vitro* transcription (IVT) reaction

**Figure S2.** Analysis of unmodified and chemically modified dsRNA synthesised by IVT.

**Figure S3.** Degradation of unmodified and chemically modified dsRNA using stink bug salival nucleases.

**Figure S4.** Colorado potato beetle (CPB) gut secretion stability assay and UV stability assay to determine the resistance of chemically modified dsRNA to degradation by insect gut secretions containing nucleases, and degradation by exposure to UV radiation.

**Figure S5.** Schematic illustration of the domain architecture and mechanism of Dicer and RNase III family enzymes.

**Figure S6.** Analysis of variation of RL values with concentration for unmodified, phosphorothioate, 2'-fluoro and 5-hydroxymethyl dsRNA.

**Figure S7.** Comparison of reproducibility of luciferase assay results for unmodified FLuc dsRNA.

**Figure S8.** Extended data for i*n vitro* analysis of the effects of chemically modified dsRNA modifications on RNAi in insect cells across a range of dsRNA concentrations.

**Figure S9.** WCR survival feeding assay using Target B scrambled control dsRNA.

**Figure S10.** WCR survival feeding assay using chemically modified dsRNA (1).

**Figure S11.** WCR survival feeding assay using chemically modified dsRNA (1) , Day 7 final percentage survival.

**Figure S12.** WCR survival feeding assay using chemically modified dsRNA (2).

**Figure S13.** WCR modified 1PS dsRNA plate feeding assay.

**Figure S14.** WCR modified 2PS dsRNA plate feeding assay.

**Figure S15.** WCR modified 1&2PS dsRNA soil feeding assay, week 0 time point.

**Figure S16.** WCR modified 1&2PS dsRNA soil feeding assay, week 0 time point, Day 7 final percentage survival.

**Table S1.**

Primers used in the PCR synthesis of DNA templates for IVT reactions. T7 polymerase binding sites are marked in bold.

| | |
|---|---|
| FLuc – Forward | 5’-**TAATACGACTCACTATAG**GGTGGCGCCCCTAGATG-3’ |
| FLuc – Reverse | 5’-**TAATACGACTCACTATAG**GGCGACGCCCGCTGATA-3’ |
| F59C6.5 – Forward | 5’-**TAATACGACTCACTATAG**GGTGGCGCCCCTAGATG-3’ |
| F59C6.5 – Reverse | 5’-**TAATACGACTCACTATAG**GGCGACGCCCGCTGATA -3’ |
| GFP Passenger (Sense) Strand – Forward | 5’-GCG**TAATACGACTCACTATAG**GAGATACCCAGATCATATGAAACGG-3’ |
| GFP Passenger (Sense) Strand – Reverse | 5’-CAATTTGTGTCCAAGAATGTTTCC-3’ |
| GFP Guide (Antisense) Strand – Forward | 5’-AGATACCCAGATCATATGAAACGG-3’ |
| GFP Guide (Antisense) Strand – Reverse | 5’-GCG**TAATACGACTCACTATAG**GCAATTTGTGTCCAAGAATGTTTCC-3’ |

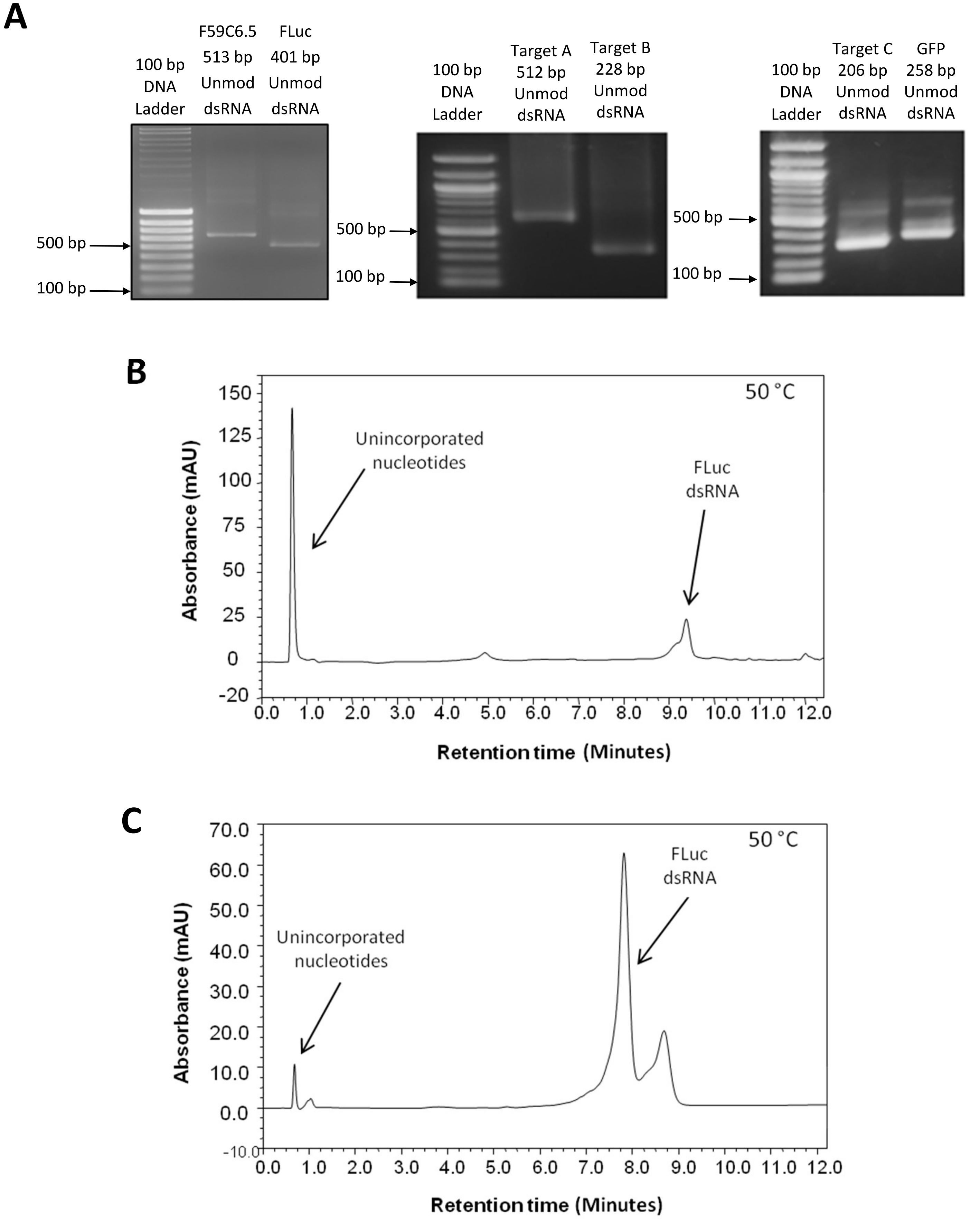


**Figure S1. Analysis of dsRNA synthesised by *in vitro* transcription (IVT) reaction**

(**A**) Gel electrophoretogram of unmodified Target A, Target B, Target C, GFP, F59C6.5, and FLuc dsRNAs synthesised by *in vitro* transcription (IVT) using T7 RNA polymerase. (**B, C**) Ion pair reverse phase HPLC analysis of unmodified dsRNA in non-denaturing conditions at 50 °C. **(B)** Chromatogram of unpurified FLuc dsRNA generated via IVT. (**C**) Chromatogram of purified FLuc dsRNA, purified using solid phase extraction (SPE), demonstrating the removal of unincorporated NTP contaminants.

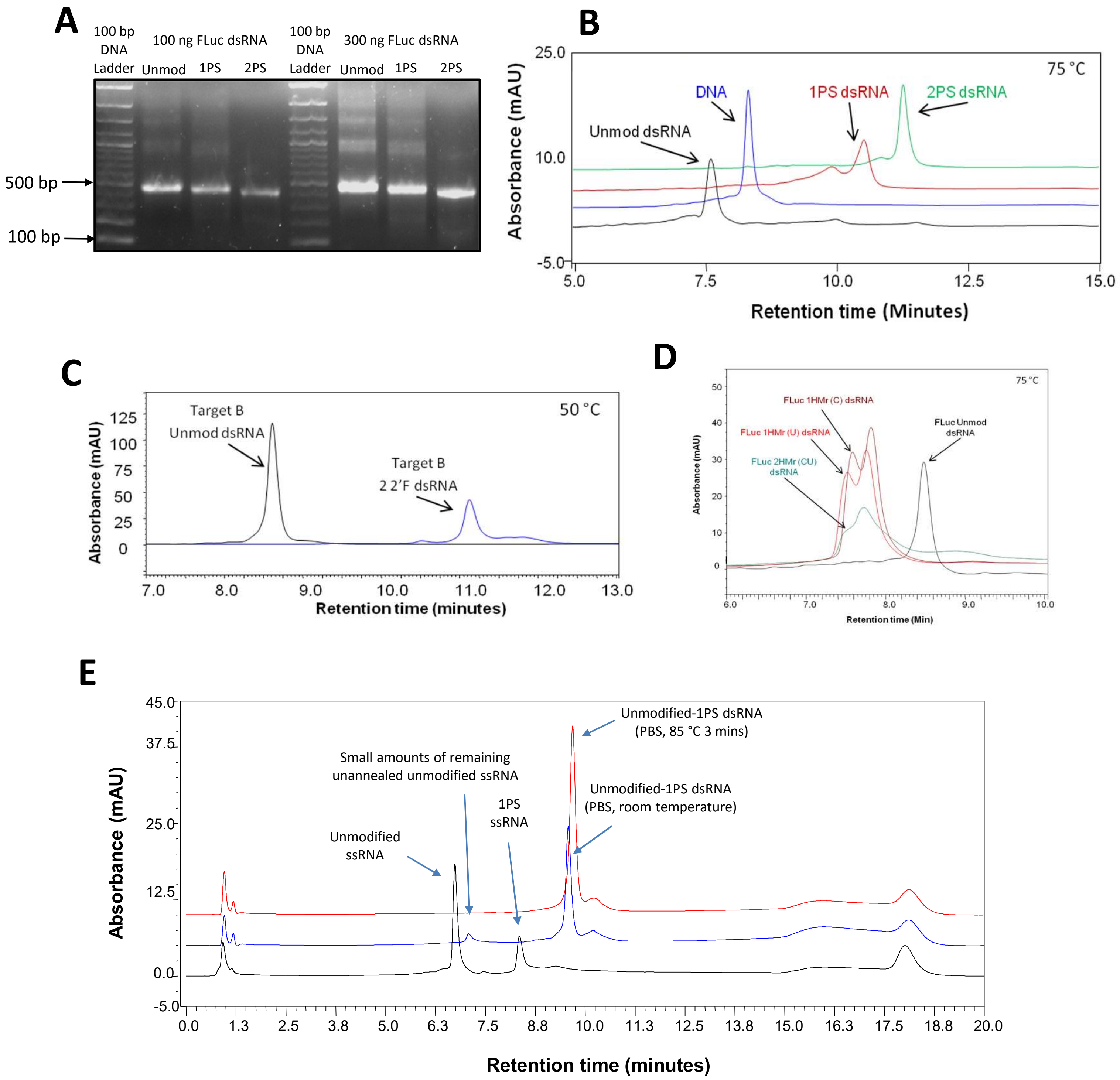


**Figure S2. Analysis of unmodified and chemically modified dsRNA synthesised by IVT.**

Validation of the incorporation of the phosphorothioate modifications in IVT synthesised RNA was confirmed by gel electrophoresis analysis and IP RP HPLC. A) Gel electrophoretogram of 100 ng and 300 ng of unmodified, 1PS and 2PS FLuc dsRNA. A shift in electrophoretic mobility of phosphorothioate dsRNA compared to unmodified dsRNA is observed. B) IP RP HPLC analysis of 1PS and 2PS dsRNA under denaturing conditions at 75 °C using gradient 1 demonstrated an increase in retention time as a result of increasing hydrophobicity of the phosphorothioate modified RNA. C/D) IP RP HPLC analysis of 2'-fluoro and 5-hydroxymethyl modified RNA. Validation of the incorporation of the 2'-fluoro and 5-hydroxymethyl modifications in IVT synthesised RNA was also confirmed by IP RP HPLC analysis, which demonstrated retention time shifts due to changes in hydrophobicity due to the presence of chemical modifications in the RNA. E) Analysis and validation of the annealing of ssRNA to form duplex dsRNA. IP RP HPLC analysis was also utilised in order to confirm full annealing of complimentary ssRNAs to dsRNA, and that no ssRNA contaminants remained . The results show that annealing of unmodified ssRNA and its complementary 1PS ssRNA anneal fully at room temperature in PBS, leaving small quantities of any ssRNA that was present in excess. Heating the sample in 1X PBS to 85°C for 3 minutes followed by cooling to room temperature prior to analysis, results in a similar yield of dsRNA and degrades any remaining excess ssRNA.

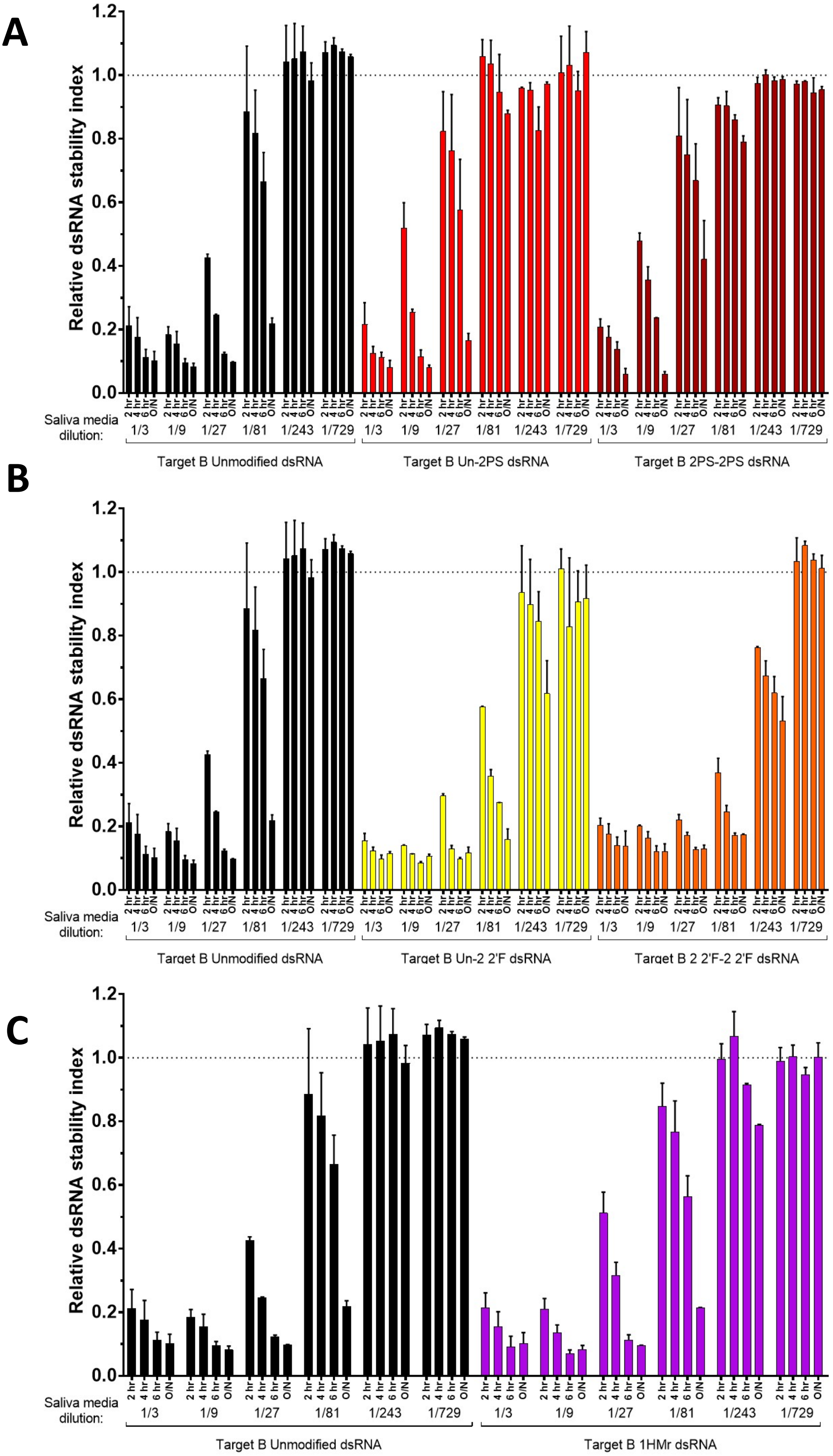


**Figure S3. Degradation of unmodified and chemically modified dsRNA using stink bug salival nucleases.**

Bar graphs quantifying degradation of unmodified and chemically modified dsRNA analysed by gel electrophoresis with gel band intensity quantified in ImageJ. dsRNA incubated at room temperature with Sf9 cell culture medium containing stink bug saliva. Saliva contaminated medium at a range of dilutions in water, and reactions stopped at either 2 hrs , 4 hrs, 6hrs or after incubation overnight (O/N) by addition of formamide loading dye and freezing at -20 °C. (A) Bar graph showing quantification of gel band intensity for unmodified and phosphorothioate dsRNA. (B) Bar graph showing quantification of gel band intensity for unmodified and 2'-fluoro dsRNA. (C) Bar graph showing quantification of gel band intensity for unmodified and 5-hydroxymethyl dsRNA. For all graphs the mean and SD of two replicates are shown.

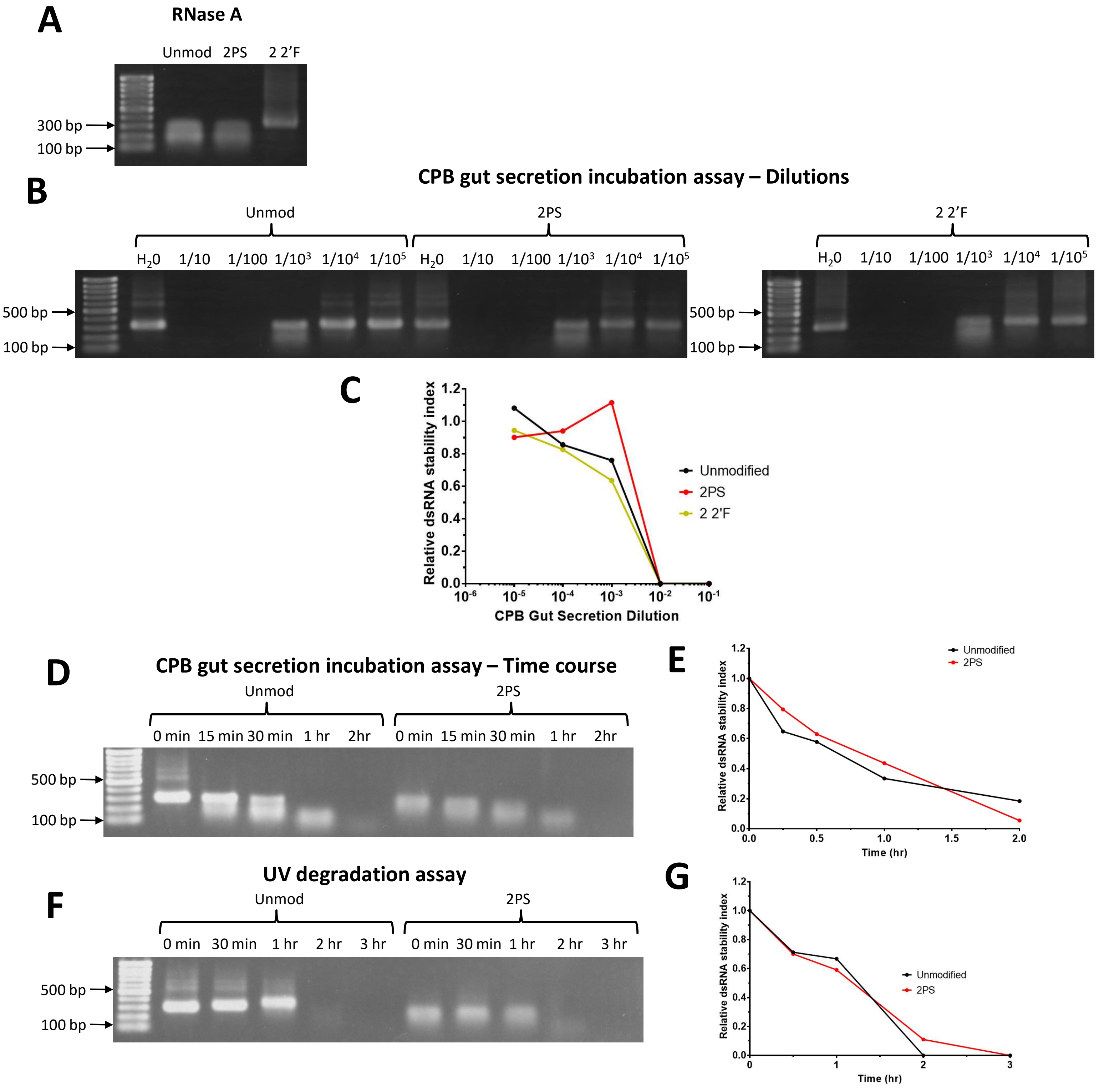


**Figure S4. Colorado potato beetle (CPB) gut secretion stability assay and UV stability assay to determine the resistance of chemically modified dsRNA to degradation by insect gut secretions containing nucleases, and degradation by exposure to UV radiation.**

(**A**) Effect of RNase A incubation on unmodified, phosphorothioate and 2'-fluoro dsRNA. RNase A incubated with dsRNA for 20 mins at 37 °C followed by addition of formamide loading dye and immediate gel electrophoresis analysis. (**B/C**) CPB gut secretion stability assay. CPB gut secretions diluted in water, and incubated with dsRNA for 30 mins at 37 °C. Reactions stopped by addition of formamide loading dye and freezing at -20 °C. Reactions were then analysed by gel electrophoresis and quantified using ImageJ. (**B**) Gel electrophoretograms of unmodified, 2PS, and 2 2'F Target B dsRNA incubated with CPB gut secretion dilutions. (**C**) Graph showing quantification of gel band intensity from (**B**). (**D/E**) CPB gut secretion stability assay: time course. CPB gut secretions diluted in water to 1/1,000 and incubated with dsRNA at 37 °C. Samples removed at 15 min, 30 min, 1hr and 2hr time points. Reactions stopped by addition of formamide loading dye and freezing at -20 °C. Reactions were then analysed by gel electrophoresis and quantified using ImageJ. (**D**) Gel electrophoretograms of unmodified and 2PS Target B dsRNA incubated with CPB gut secretion dilution. (**E**) Time course graph showing quantification of gel band intensity from (**D**).

(**F, G**) UV degradation assay. Aqueous dsRNA solutions in UV-permeable microcuvettes were placed in a UV crosslinker and exposed to UV radiation, with samples removed at 30 min, 1hr, 2hr and 3hr time points and stored at -20 °C, followed by gel electrophoresis analysis and quantified using ImageJ. (**F**) Gel electrophoretograms of unmodified and 2PS Target B dsRNA exposed to UV radiation. (**G**) Time course graph showing quantification of gel band intensity from (**F**).

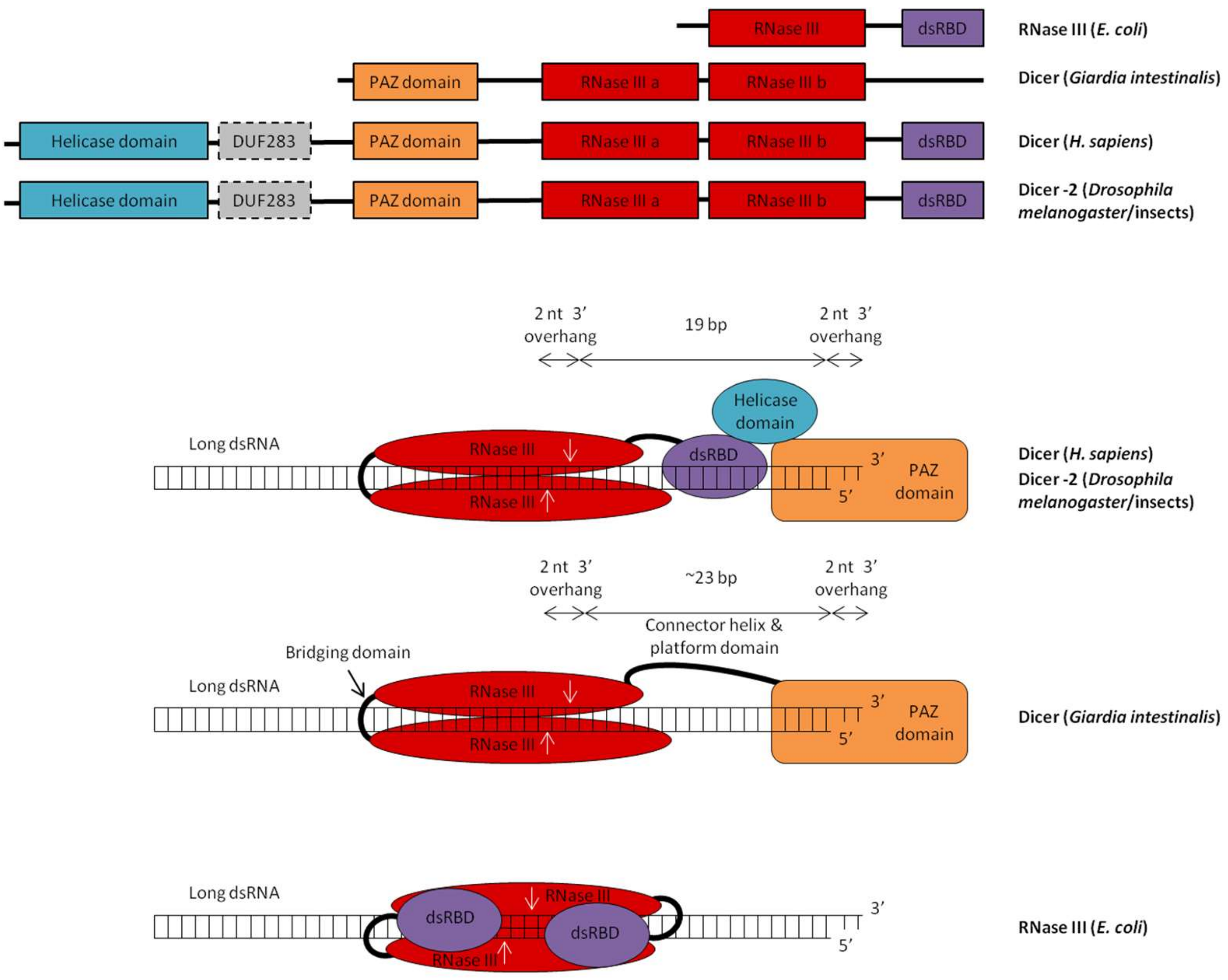


**Figure S5. Schematic illustration of the domain architecture and mechanism of Dicer and RNase III family enzymes.**

Top: Domain architecture of bacterial, *Giardia*, human and insect Dicer/RNase III family enzymes. DUF283 (Domain of unknown function 283) is known to be an atypical dsRNA binding domain. Human, *Drosophila* and other insect Dicers have similar domain architecture, however the size and sequence of the main functional domains, linking domains, and overall enzyme are different for each (domain sizes not to scale). Bottom: Cleavage mechanism of human/insect, *Giardia*, and bacterial Dicer/RNase III family enzymes. Dicer enzymes form internal psuedo-dimers between the RNase III domains to cut both dsRNA strands, whereas two separate bacterial RNase III monomers form a true dimer to cut both strands. Cut sites indicated by white arrows. Not to scale.

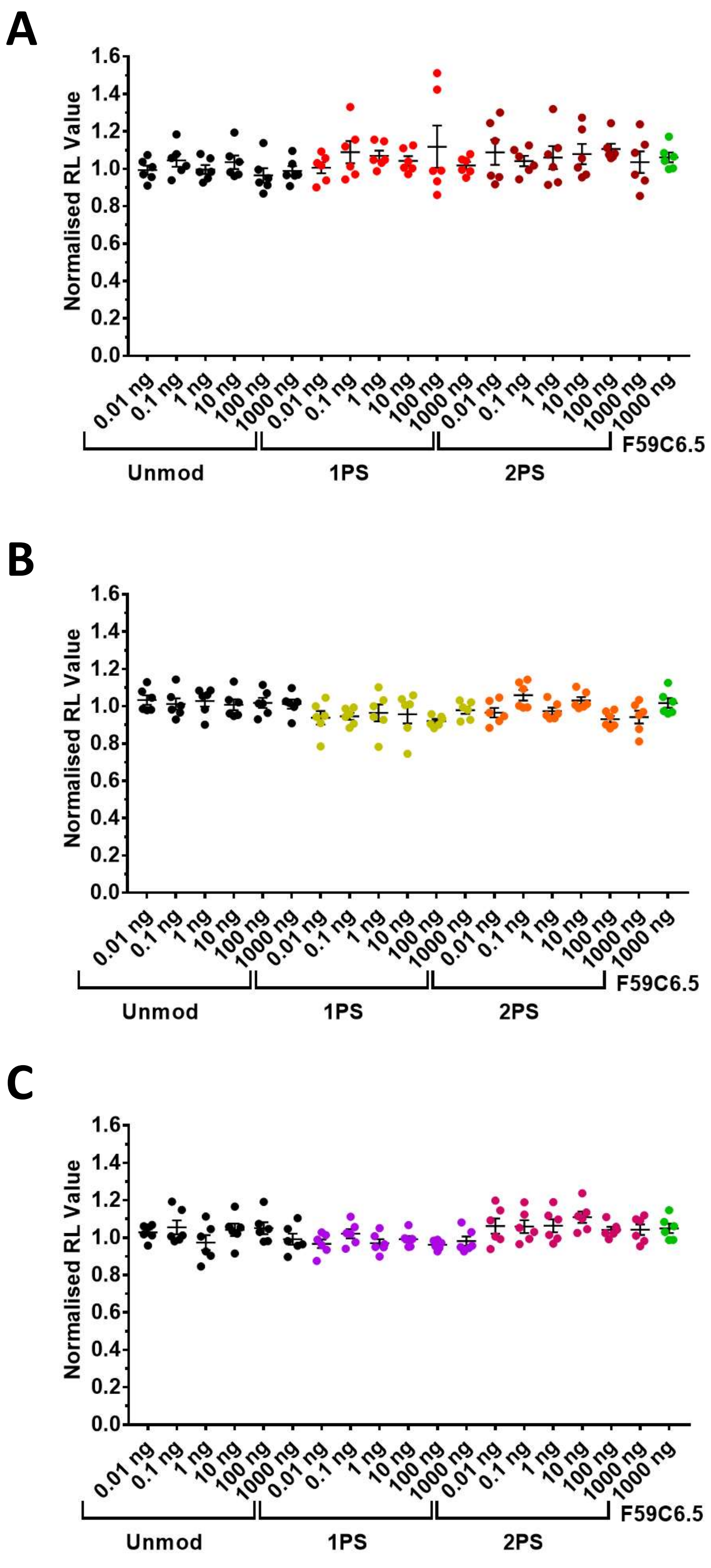


**Figure S6. Analysis of variation of RL values with concentration for unmodified, phosphorothioate, 2'-fluoro and 5-hydroxymethyl dsRNA.** Graphs of normalised RL values from the luciferase assay results for all concentrations of PS (**A**), 2'F (**B**), and HMr (**C**) FLuc dsRNAs compared to the unmodified FLuc and unmodified F59C6.5 controls. *Renilla* luciferase luminescence intensity (RL) normalised against RL values for control conditions with no dsRNA. n = 6.

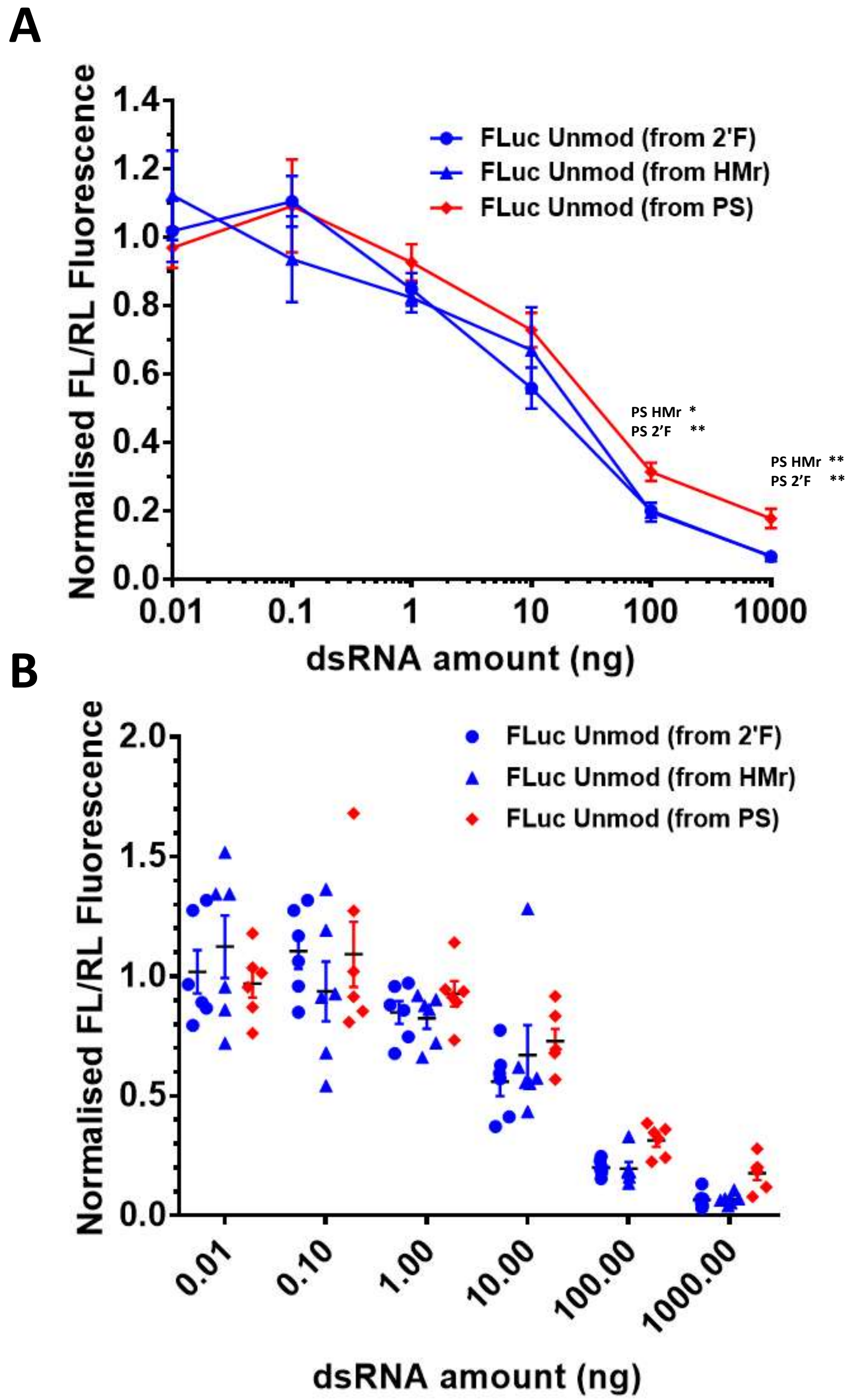


**Figure S7. Comparison of reproducibility of luciferase assay results for unmodified FLuc dsRNA.**
Comparison of reproducibility of luciferase assay results for unmodified (Unmod) FLuc dsRNA between the luciferase assays screening 2’F and HMr dsRNA (conducted concurrently), and PS dsRNA (conducted separately). Results showing quantification of RNAi effects on a firefly luciferase reporter in *Drosophila* Kc167 cell cultures, quantified using a dual luciferase assay reporter system. RNAi effects on a firefly luciferase reporter by FLuc dsRNA, is presented as ratios (FL/RL) of firefly luciferase luminescence intensity (FL) to control *Renilla* luciferase luminescence intensity (RL), normalised against FL/RL values for control conditions with no dsRNA. Greater loss of fluorescence correlates to greater levels of RNAi effect. (**A**) Dose curves of normalised FL/RL values plotted against log of dsRNA dose per well in ng. (**B**) Individual normalised FL/RL values for the dose curves in figure a.
In all graphs, mean and SEM are plotted (n=6). Stars denote significance as calculated by unpaired T-tests. ns = P > 0.05, * = P ≤ 0.05, ** = P ≤ 0.01, *** = P ≤ 0.001, **** = P ≤ 0.0001.

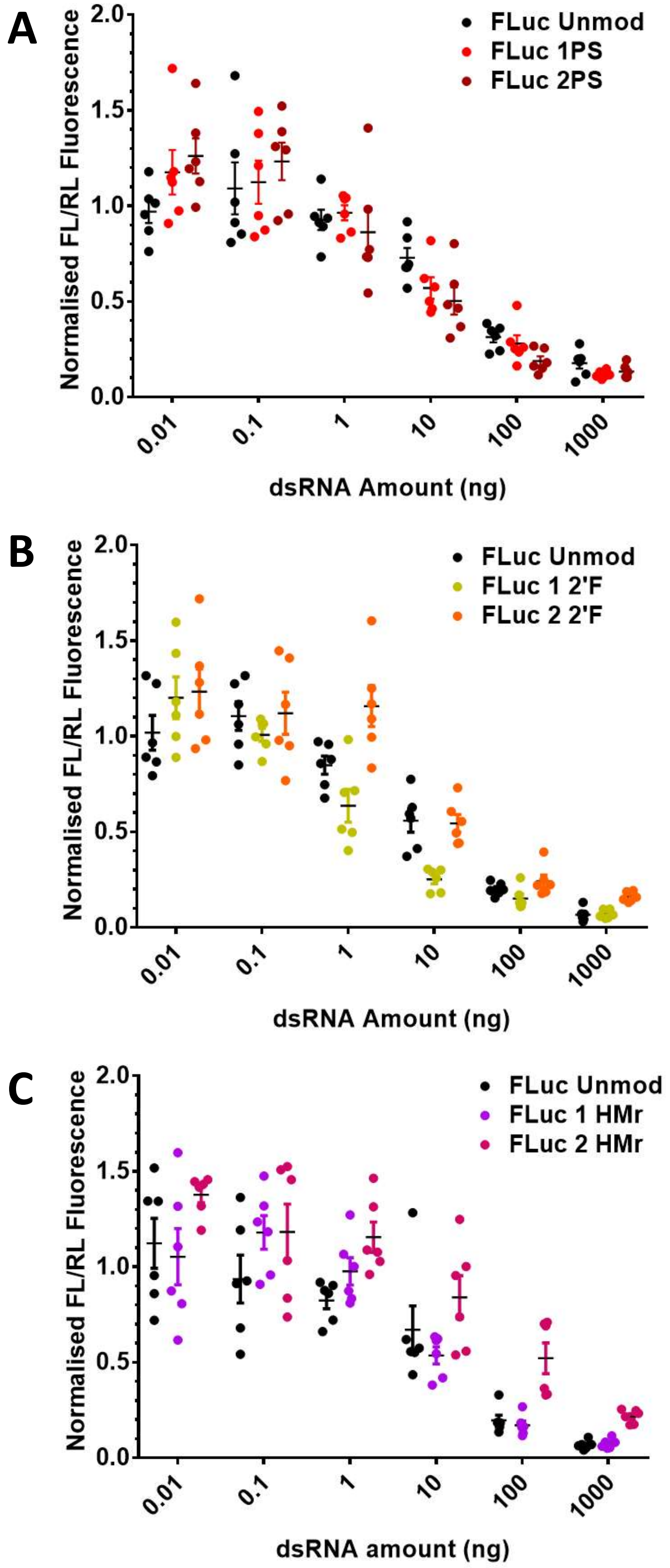


**Figure S8. Extended data for i*n vitro* analysis of the effects of chemically modified dsRNA modifications on RNAi in insect cells across a range of dsRNA concentrations.**

Results showing quantification of RNAi effects on a firefly luciferase reporter in *Drosophila* Kc167 cell cultures, quantified using a dual luciferase assay reporter system. RNAi effect on a firefly luciferase reporter by FLuc dsRNA, is presented as ratios (FL/RL) of firefly luciferase luminescence intensity (FL) to control *Renilla* luciferase luminescence intensity (RL), normalised against FL/RL values for control conditions with no dsRNA. Graphs show individual normalised FL/RL values for the dose curves in **Figure 4 C, F, I**. (**A**) Results for phosphorothioate (PS) and unmodified (Unmod) FLuc dsRNA. (**B**) Results for 2′-fluoro (2′F) and unmodified (Unmod) FLuc dsRNA. (**C**) Results for 5-hydroxymethyl (HMr) and unmodified (Unmod) FLuc dsRNA.

In all graphs, mean and SEM are plotted (n=6). Stars denote significance as calculated by unpaired T-tests. ns = P > 0.05, * = P ≤ 0.05, ** = P ≤ 0.01, *** = P ≤ 0.001, **** = P ≤ 0.0001.

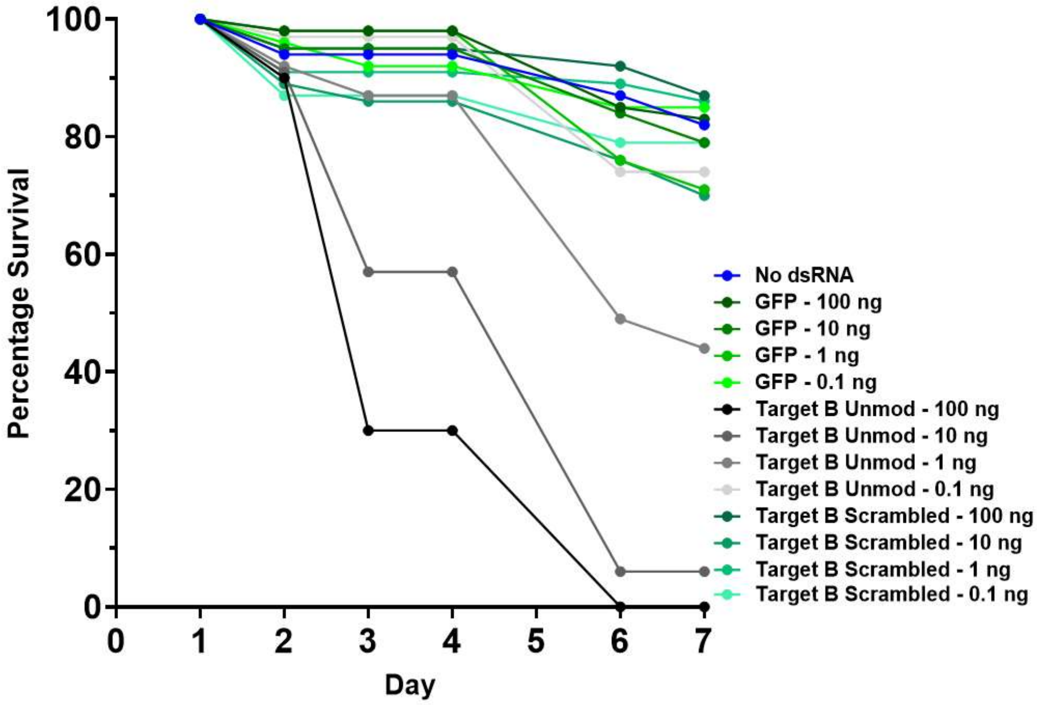


**Figure S9. WCR survival feeding assay using Target B scrambled control dsRNA.**
WCR plate feeding assay. WCR were fed on an artificial diet containing 0.1 to 100 ng of dsRNA in well plates and mortality measured over 7 days. Unmod dsRNA n = 55, 53, 52, 46, GFP dsRNA n = 51, 51, 53, 56, Scrambled dsRNA n = 48, 54, 51, 54, No dsRNA n = 114. Survival timecourse shown. Day 7 survival data in Figure 7B.

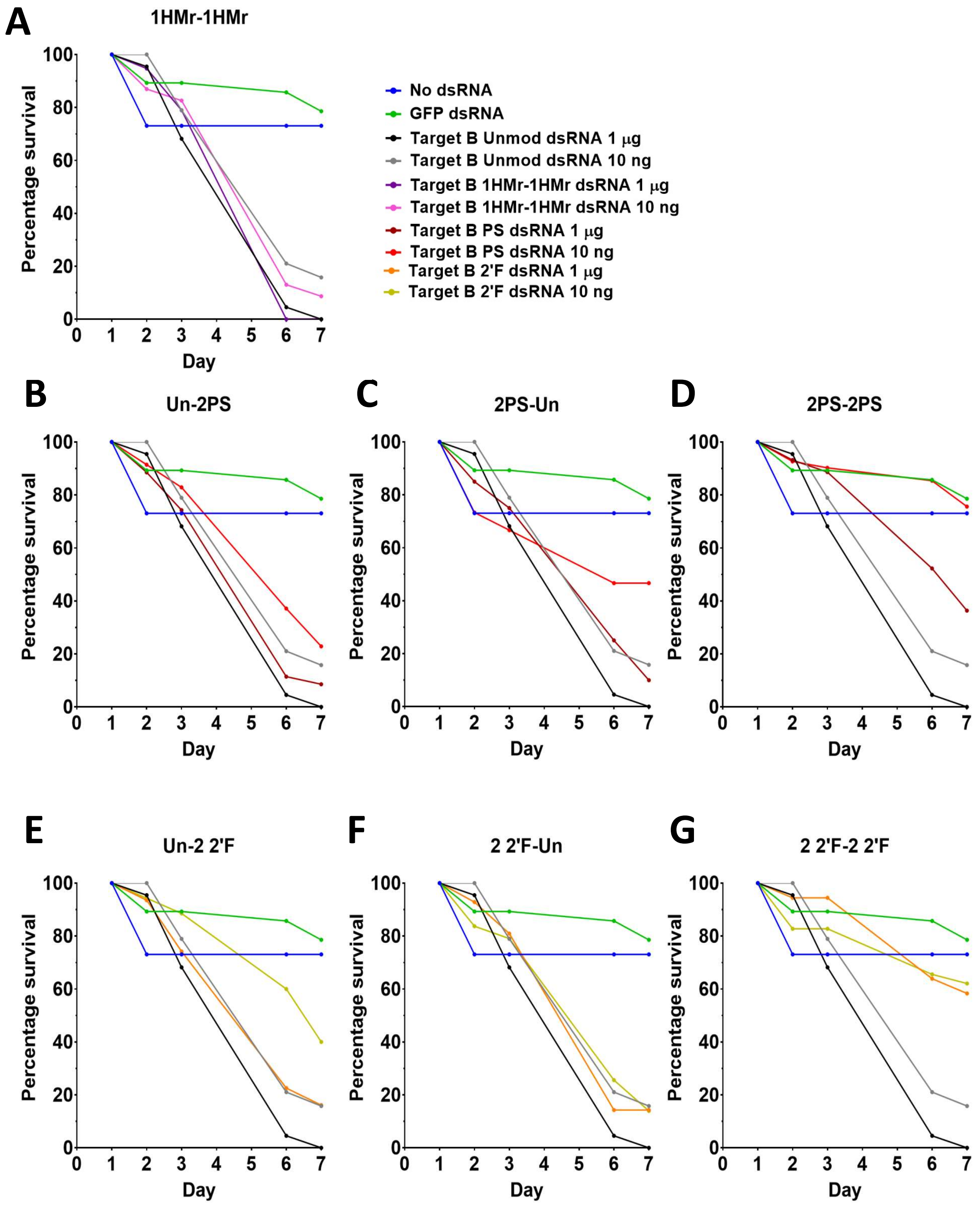


**Figure S10. WCR survival feeding assay using chemically modified dsRNA (1).**

(**A**) WCR modified HMr dsRNA plate feeding assay survival timecourse. WCR fed on an artificial diet containing dsRNA in well plates and mortality measured over 7 days. GFP n = 46, Unmod n = 46, 45, 1HMr-1HMr n = 45, 38, No dsRNA n = 48. (**B-D**) WCR modified PS dsRNA plate feeding assay survival timecourses. WCR fed on an artificial diet containing dsRNA in well plates and mortality measured over 7 days. GFP n = 46, Unmod n = 46, 45, Un-2PS n = 44, 45, 2PS-Un n = 45, 43, 2PS-2PS n = 47, 48, No dsRNA n = 48. (**B**) Survival timecourse, for Un-2PS dsRNA. (**C**) Survival timecourse for 2PS-Un dsRNA. (**D**) Survival timecourse for 2PS-2PS dsRNA. (**E-G**) WCR modified 2’F dsRNA plate feeding assay. WCR fed on an artificial diet containing dsRNA in well plates and mortality measured over 7 days. GFP n = 46, Unmod n = 46, 45, Un-2 2’F n = 42, 44, 2 2’F-Un n = 44, 50, 2 2’F-2 2’F n = 45, 48, No dsRNA n = 48. (**E**) Survival timecourse, for Un-2 2’F dsRNA. (**F**) Survival timecourse for 2 2’F-Un dsRNA. (**G**) Survival timecourse for 2 2’F-2 2’F dsRNA.

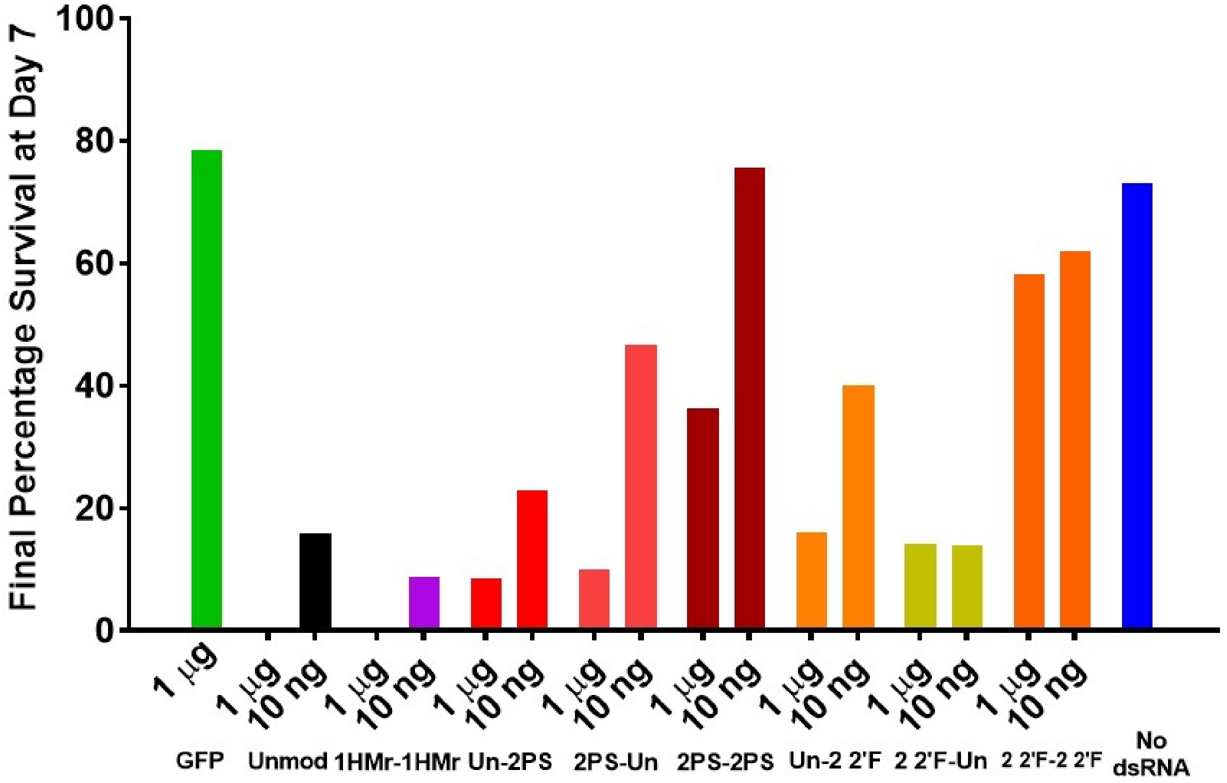


**Figure S11. WCR survival feeding assay using chemically modified dsRNA (1) , Day 7 final percentage survival.**
WCR chemically modified dsRNA plate feeding assay survival data for day 7 of the timecourse assay data in Figure S9. Number of insects used for each dsRNA concentration (L-R): GFP n = 46, Unmod n = 46, 45, 1HMr-1HMr n = 45, 38, Un-2PS n = 44, 45, 2PS-Un n = 45, 43, 2PS-2PS n = 47, 48, Un-2 2’F n = 42, 44, 2 2’F-Un n = 44, 50, 2 2’F-2 2’F n = 45, 48, No dsRNA n = 48.

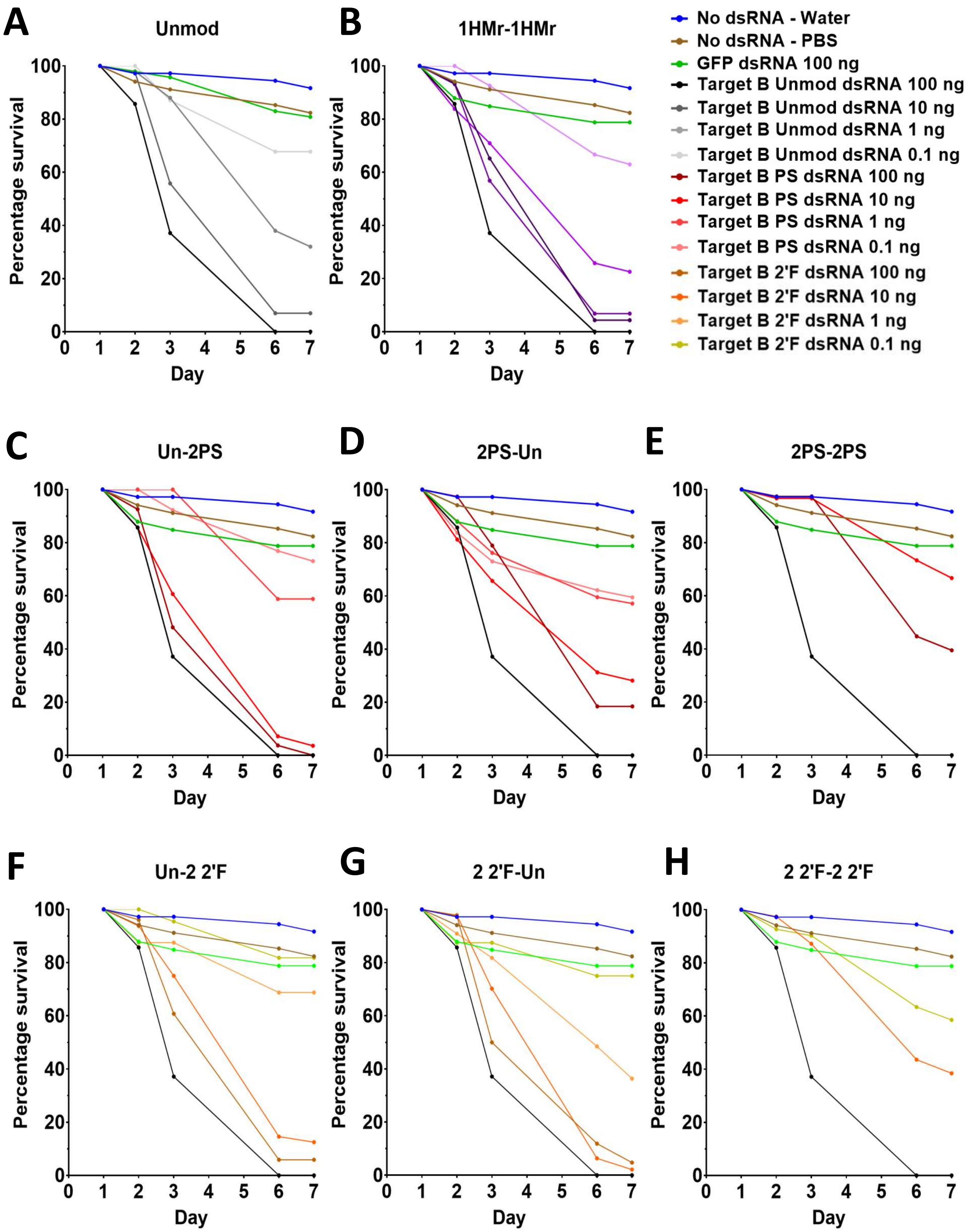


**Figure S12. WCR survival feeding assay using chemically modified dsRNA (2).**

(**A**) WCR unmodified dsRNA plate feeding assay survival timecourse. WCR fed on an artificial diet containing dsRNA in well plates and mortality measured over 7 days. GFP n = 50, Unmod n = 47, 48, 54, 45, No dsRNA - Water n = 46, No dsRNA – PBS n = 38. (**B**) WCR modified HMr dsRNA plate feeding assay survival timecourse. WCR fed on an artificial diet containing dsRNA in well plates and mortality measured over 7 days. Survival timecourse for HMr dsRNA. GFP n = 50, Unmod n = 47, 1HMr-1HMr n = 53, 58, 46, 41, No dsRNA - Water n = 46, No dsRNA – PBS n = 38. (**C-E**) WCR modified PS dsRNA plate feeding assay. WCR fed on an artificial diet containing dsRNA in well plates and mortality measured over 7 days. GFP n = 50, Unmod n = 47, Un-2PS n = 49, 48, 51, 62, 2PS-Un n = 51, 52, 54, 60, 2PS-2PS n = 48, 36, No dsRNA - Water n = 46, No dsRNA – PBS n = 38. (**C**) Survival timecourse, for Un-2PS dsRNA. (**D**) Survival timecourse for 2PS-Un dsRNA. (**E**) Survival timecourse for 2PS-2PS dsRNA. (**F-H**) WCR modified 2’F dsRNA plate feeding assay. WCR fed on an artificial diet containing dsRNA in well plates and mortality measured over 7 days. GFP n = 50, Unmod n = 47, Un-2 2’F n = 52, 66, 38, 43, 2 2’F-Un n = 49, 58, 47, 48, 2 2’F-2 2’F n = 41, 46, No dsRNA - Water n = 46, No dsRNA – PBS n = 38. (**F**) Survival timecourse, for Un-2 2’F dsRNA. (**G**) Survival timecourse for 2 2’F-Un dsRNA. (**H**) Survival timecourse for 2 2’F-2 2’F dsRNA.

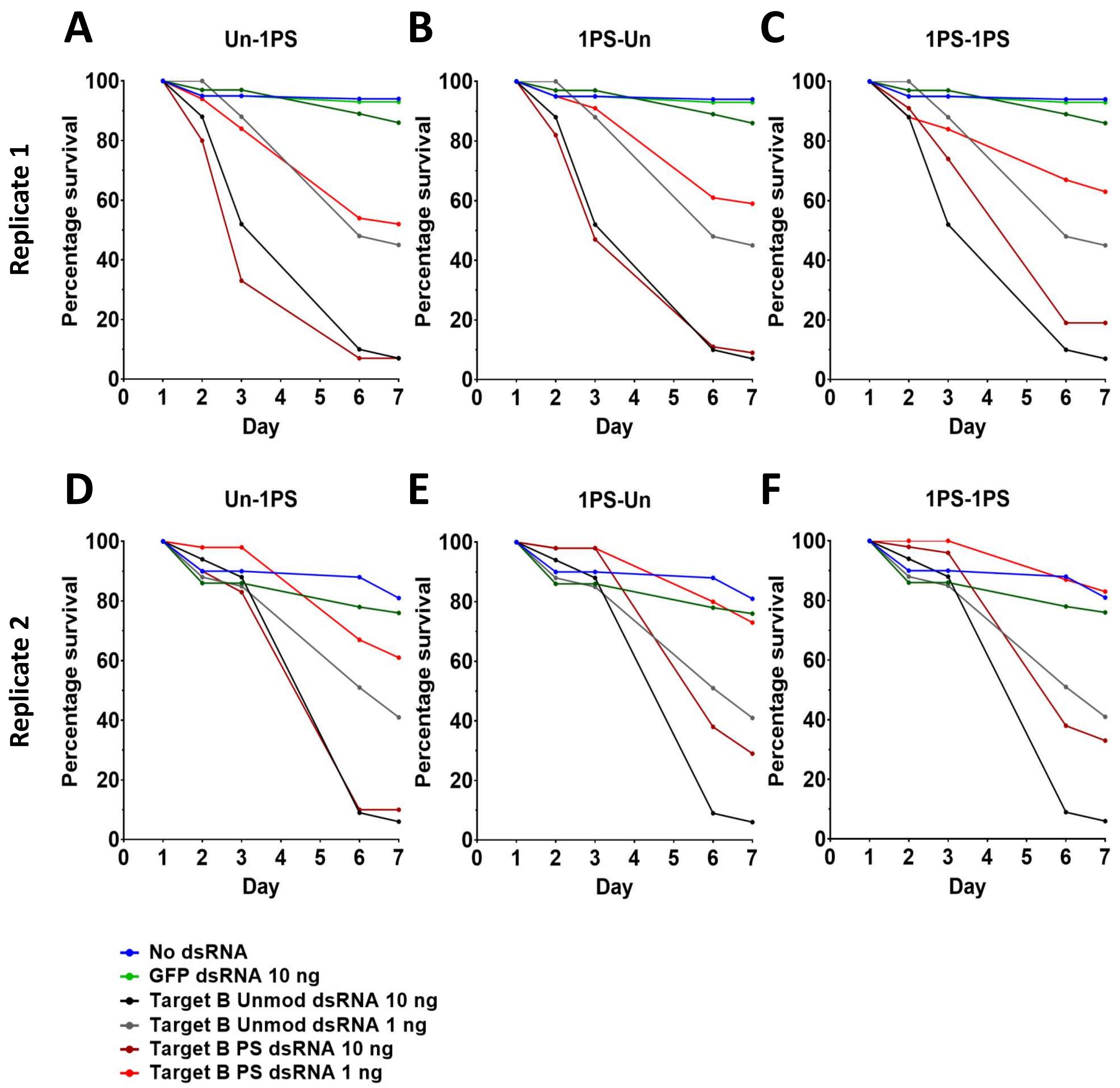


**Figure S13. WCR modified 1PS dsRNA plate feeding assay.**

WCR fed on an artificial diet containing dsRNA in well plates and mortality measured over 7 days. GFP Rep 1 n = 47, Rep 2 n = 50; Unmod Rep 1 n = 47, 43, Rep 2 n = 49, 54; Un-1PS Rep 1 n = 45, 52, Rep 2 n = 47, 49; 1PS-Un Rep 1 n = 48, 48, Rep 2 n = 46, 48; 1PS-1PS Rep 1 n = 48, 49, Rep 2 n = 50, 48; No dsRNA Rep 1 n = 96, Rep 2 n = 97; (**A**) Survival timecourse for Un-1PS dsRNA, replicate 1. (**B**) Survival timecourse, for 1PS-Un dsRNA, replicate 1. (**C**) Survival timecourse for 1PS-1PS dsRNA, replicate 1. (**D**) Survival timecourse for Un-1PS dsRNA, replicate 2. (**E**) Survival timecourse for 1PS-Un dsRNA, replicate 2. (**F**) Survival timecourse for 1PS-1PS dsRNA, replicate 2.

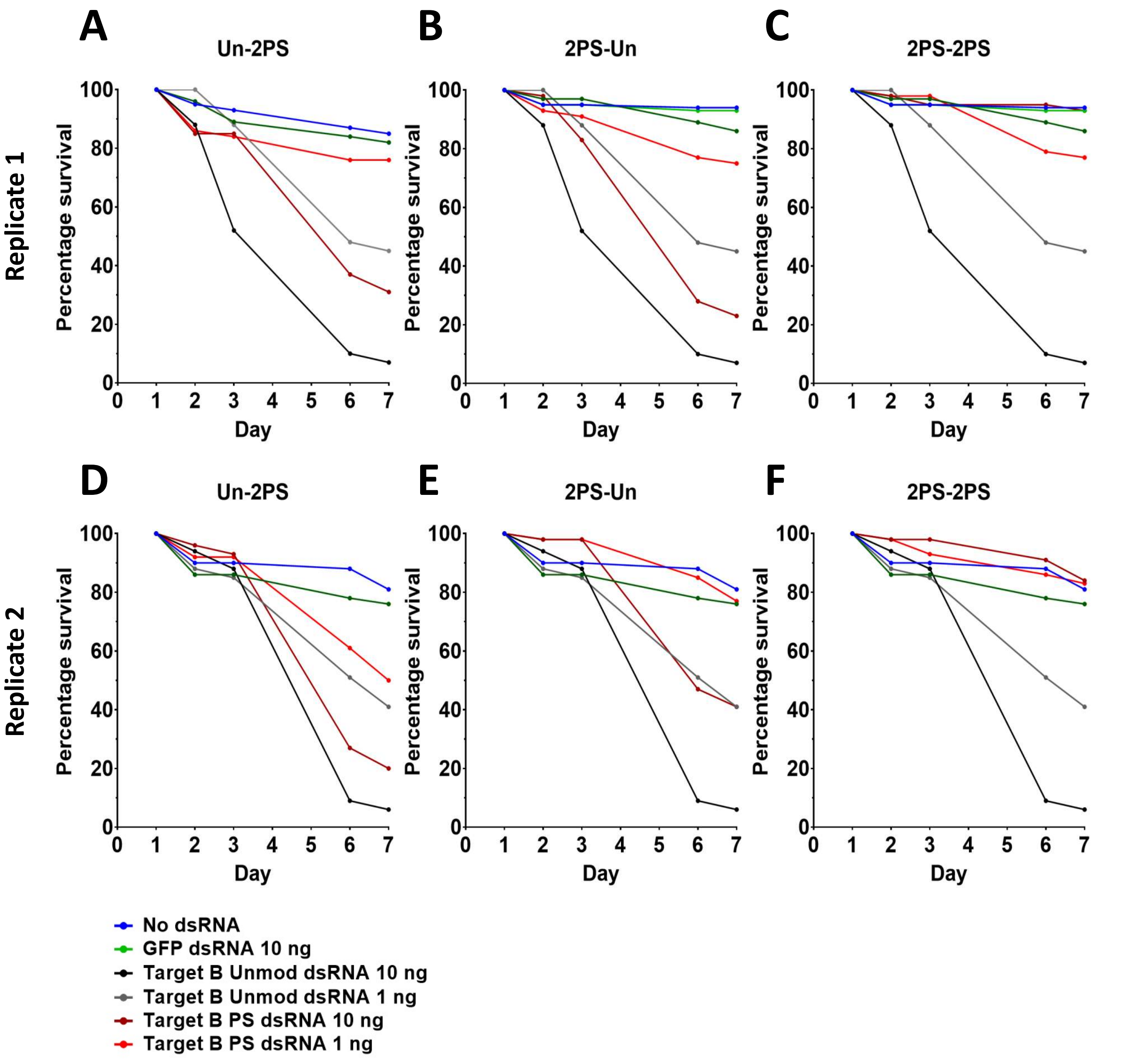


**Figure S14. WCR modified 2PS dsRNA plate feeding assay.**

WCR fed on an artificial diet containing dsRNA in well plates and mortality measured over 7 days. GFP Rep 1 n = 47, Rep 2 n = 50; Unmod Rep 1 n = 47, 43, Rep 2 n = 49, 54; Un-2PS Rep 1 n = 55, 54, Rep 2 n = 50, 42; 2PS-Un Rep 1 n = 44, 50, Rep 2 n = 49, 48; 2PS-2PS Rep 1 n = 49, 48, Rep 2 n = 46, 45; No dsRNA Rep 1 n = 96, Rep 2 n = 97; (**A**) Survival timecourse for Un-2PS dsRNA, replicate 1. (**B**) Survival timecourse, for 2PS-Un dsRNA, replicate 1. (**C**) Survival timecourse for 2PS-2PS dsRNA, replicate 1. (**D**) Survival timecourse for Un-2PS dsRNA, replicate 2. (**E**) Survival timecourse for 2PS-Un dsRNA, replicate 2. (**F**) Survival timecourse for 2PS-2PS dsRNA, replicate 2.

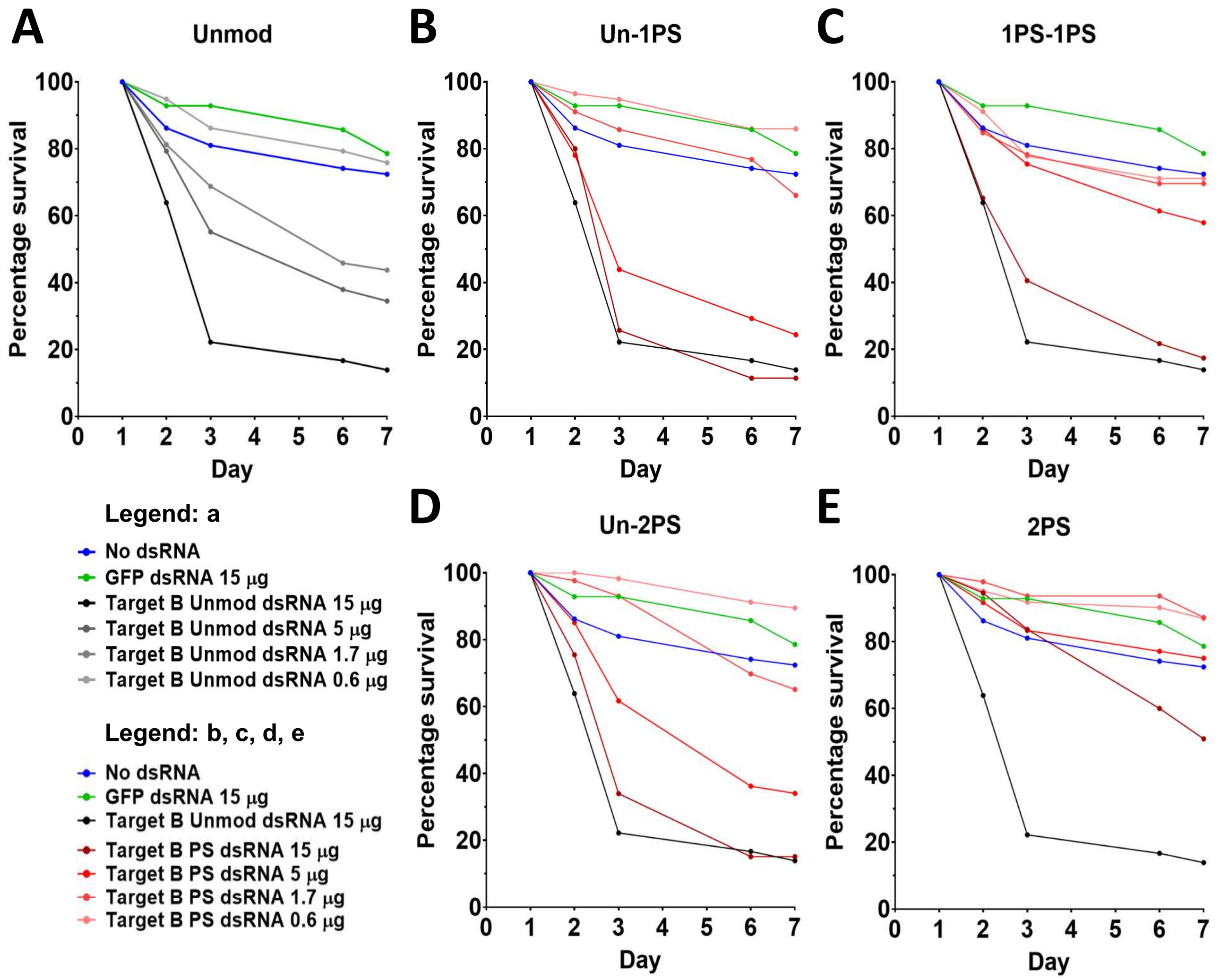


**Figure S15. WCR modified 1&2PS dsRNA soil feeding assay, week 0 time point.**

WCR were left on soil containing dsRNA for 1 day, then transferred to diet plates and mortality measured until day 7. GFP n = 124; Unmod n = 112, 126, 115; Un-1PS n = 109, 121, 124; 1PS-1PS n = 116, 121, 117; Un-2PS n = 120, 126, 117; 2PS-2PS n = 118, 125, 117; No dsRNA n = 119. (**A**) Survival timecourse for Unmod dsRNA. (**B**) Survival timecourse for Un-1PS dsRNA. (**C**) Survival timecourse for 1PS-1PS dsRNA. (**D**) Survival timecourse for Un-2PS dsRNA. (**E**) Survival timecourse for 2PS-2PS dsRNA.

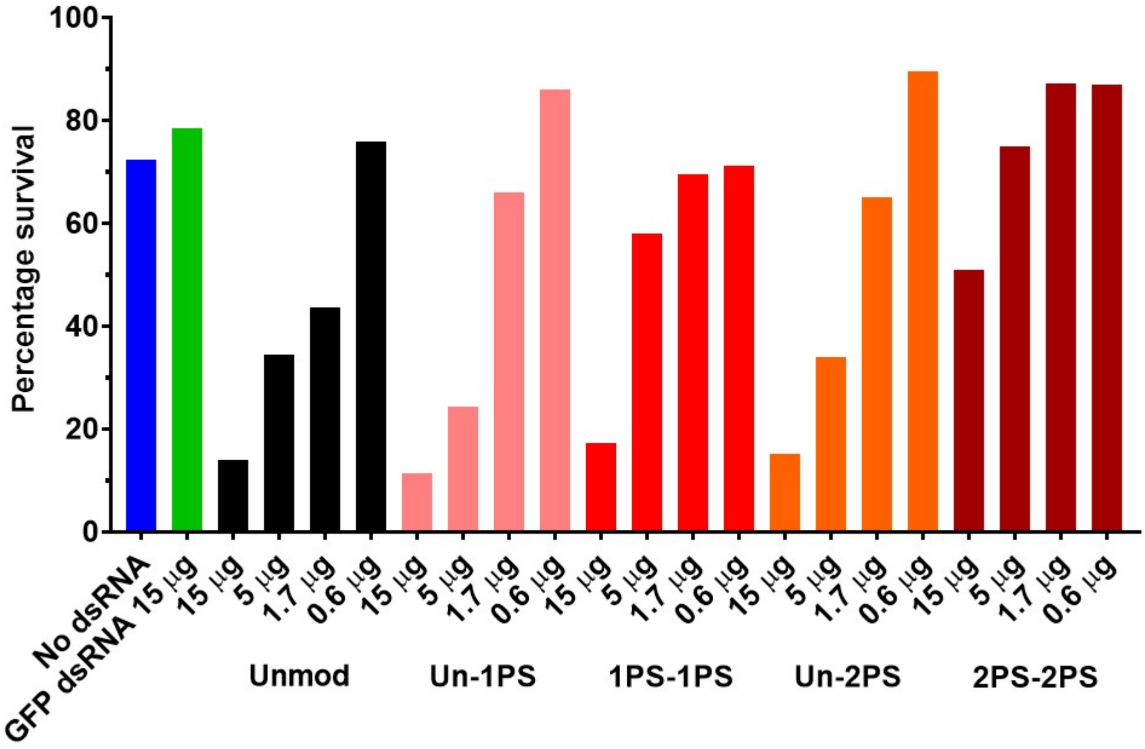


**Figure S16. WCR modified 1&2PS dsRNA soil feeding assay, week 0 time point, Day 7 final percentage survival.**

WCR chemically modified dsRNA soil feeding assay survival data for day 7 of the week 0 timecourse assay data in Figures S13&14, showing additional dsRNA concentrations beyond those included in Figure 6D. WCR were left on soil containing dsRNA for 1 day, then transferred to untreated diet plates and mortality measured until day 7. Number of insects used for each dsRNA concentration (L-R): No dsRNA n = 102; GFP dsRNA n = 117; Unmod n = 115, 119, 121, 112; Un-1PS n = 102, 116, 118, 113; 1PS-1PS n = 125, 116, 118, 113; Un-2PS n = 124, 114, 121, 116; 2PS-2PS n = 106, 121, 106, 120.
